# Co-regulation and functional cooperativity of FOXM1 and RHNO1 bidirectional genes in ovarian cancer

**DOI:** 10.1101/630442

**Authors:** Carter J Barger, Connor Branick, Linda Chee, Mustafa Albahrani, David Klinkebiel, Ronny Drapkin, Kunle Odunsi, Lee Zou, Adam R. Karpf

## Abstract

We report that the oncogenic transcription factor *FOXM1* is arranged in a head-to-head configuration with *RHNO1*, a gene involved in the ATR/CHK1-dependent DNA replication stress (DRS) response. *FOXM1* and *RHNO1* are both amplified and upregulated in high-grade serous ovarian cancer (HGSC). *FOXM1* and *RHNO1* expression are closely associated in normal and cancer tissues, including single cells, and a bidirectional promoter (F/R-BDP) mediates balanced expression. Targeting of FOXM1 and RHNO1 in HGSC cells using shRNA, CRISPR mutagenesis, or CRISPR interference directed to the F/R-BDP reduced DNA homologous recombination repair (HR) capacity, increased DNA damage, reduced clonogenic survival, and sensitized HGSC cells to the poly-ADP ribosylase inhibitor (PARPi) olaparib. Thus, there is functional cooperativity between FOXM1 and RHNO1 in cancer cells, and combinatorial targeting of this bidirectional gene pair may be a novel cancer therapeutic strategy. More broadly, our data provide evidence that bidirectional gene units function in human cancer.

## Introduction

FOXM1 is linked to cancer hallmarks through its function as a transcription factor, activating target genes that promote oncogenic phenotypes (Halasi and Gartel, 2013). The most common oncogenic function assigned to FOXM1 is deregulation of the G2/M checkpoint to promote cellular proliferation (Costa, 2005; Laoukili et al., 2005; Wonsey and Follettie, 2005). FOXM1 also promotes HR (Khongkow et al., 2014; Maachani et al., 2016; Monteiro et al., 2013; Park et al., 2012) and chemoresistance (Carr et al., 2010; Fang et al., 2018; Kwok et al., 2010; Tassi et al., 2017; Zhao et al., 2014). Pan-cancer analyses have demonstrated increased FOXM1 expression across a variety of cancers and a robust connection to genomic instability and poor prognosis (Barger et al., 2019; Carter et al., 2006; Gentles et al., 2015; Jiang et al., 2015; Li et al., 2017). We previously reported that *FOXM1* is overexpressed in high-grade serous ovarian cancer (HGSC), with gene amplification/copy number gains, loss of function of p53 and Rb, and activation of E2F1 and cyclin E1, as key mechanisms driving increased FOXM1 expression (Barger et al., 2019; Barger et al., 2015). Surprisingly, *FOXM1* amplification, but not *FOXM1* expression, associated with decreased HGSC patient survival (Barger et al., 2015). *FOXM1* is located within the 12p13.33 HGSC amplicon, which contains over 30 genes, suggesting that FOXM1 could potentially cooperate with other genes to promote HGSC.

RHNO1 (first called C12orf32) was initially reported to be overexpressed in breast cancer cell lines, exhibit nuclear localization, and promote the survival of a breast cancer cell line (Kim et al., 2010). RHNO1 was later identified in a high-throughput screen for DNA damage repair (DDR) regulators and shown to interact with the 9-1-1 complex (prompting the name change: <u>R</u>ad9, <u>R</u>ad1, <u>H</u>us1-Interacting <u>N</u>uclear <u>O</u>rphan) to promote ATR activation (Cotta-Ramusino et al., 2011). The RHNO1 N-terminus contains a conserved APSES (Asm1p, Phd1p, Sok2p, Efg1p and StuAp) DNA binding domain, which is necessary for interaction with 9-1-1, whereas the C-terminus is required for interactions with 9-1-1 and TOPBP1 and localization to DNA damage sites (Cotta-Ramusino et al., 2011).

Here, we show that FOXM1 and RHNO1 are co-amplified and co-expressed in HGSC, and that their expression in controlled by a bidirectional promoter (named the F/R-BDP). We additionally show that FOXM1 and RHNO1 they cooperatively promote HR, cell survival, and PARPi resistance. Co-regulation and functional cooperativity between FOXM1 and RHNO1 suggests that this bidirectional gene unit is a potential therapeutic target in HGSC and/or other cancers. More generally, our data provide evidence that cooperative bidirectional gene units functionally contribute to cancer.

## Results

### The 12p13.33 amplicon contains *FOXM1* and *RHNO1*, co-expressed bidirectional genes

*FOXM1* is located at 12p13.33, a region with frequent copy number gains in HGSC and several other cancers (Barger et al., 2019; Barger et al., 2015). 12p13.33 amplification was associated with reduced overall survival in HGSC but, unexpectedly, *FOXM1* mRNA expression did not (Barger et al., 2015). The 12p13.33 amplicon includes 33 genes (**Figure S1**). To identify genes at 12p13.33 that might functionally cooperate with FOXM1, we performed expression correlation analyses. *FOXM1* expression showed the strongest correlation with *RHNO1* expression in HGSC (**Figure 1A**), pan-cancer (data not shown), and normal human tissues (**Figure S2**). In addition, *RHNO1* mRNA expression correlated with reduced HGSC survival (data not shown). Genomic analyses of *FOXM1* and *RHNO1* revealed a head-to-head arrangement with an intervening putative bidirectional promoter (BDP) (**Figure S3**). Approximately 10% of human genes are arranged in this manner, and the gene pairs are frequently co-regulated and can function in similar biochemical pathways (Trinklein et al., 2004; Wakano et al., 2012; Yang et al., 2007). The *FOXM1/RHNO1 BDP* (F/R-BDP) region contains a CpG island (CGI), a common feature of BDPs (**Figure S3**) (Antequera, 2003; Takai and Jones, 2004; Wakano et al., 2012). BDPs are often enriched with specific histone modifications H3K4me3, H3K9ac, H3K27ac, and H3K4me2 (Bornelov et al., 2015) and, in agreement, ENCODE data revealed that H3K4me3 and H3K27Ac are enriched in bimodal peaks at the F/R-BDP, suggesting correlated transcriptional activity in both directions (**Figure S3**). ENCODE data also revealed an E2F1 peak, consistent with E2F’s known involvement in BDP regulation and prior data identifying E2F1 as a regulator of *FOXM1* expression (**Figure S3**) (Barger et al., 2019; Barger et al., 2015; Chen et al., 2014; Millour et al., 2011).

**Figure 1.**
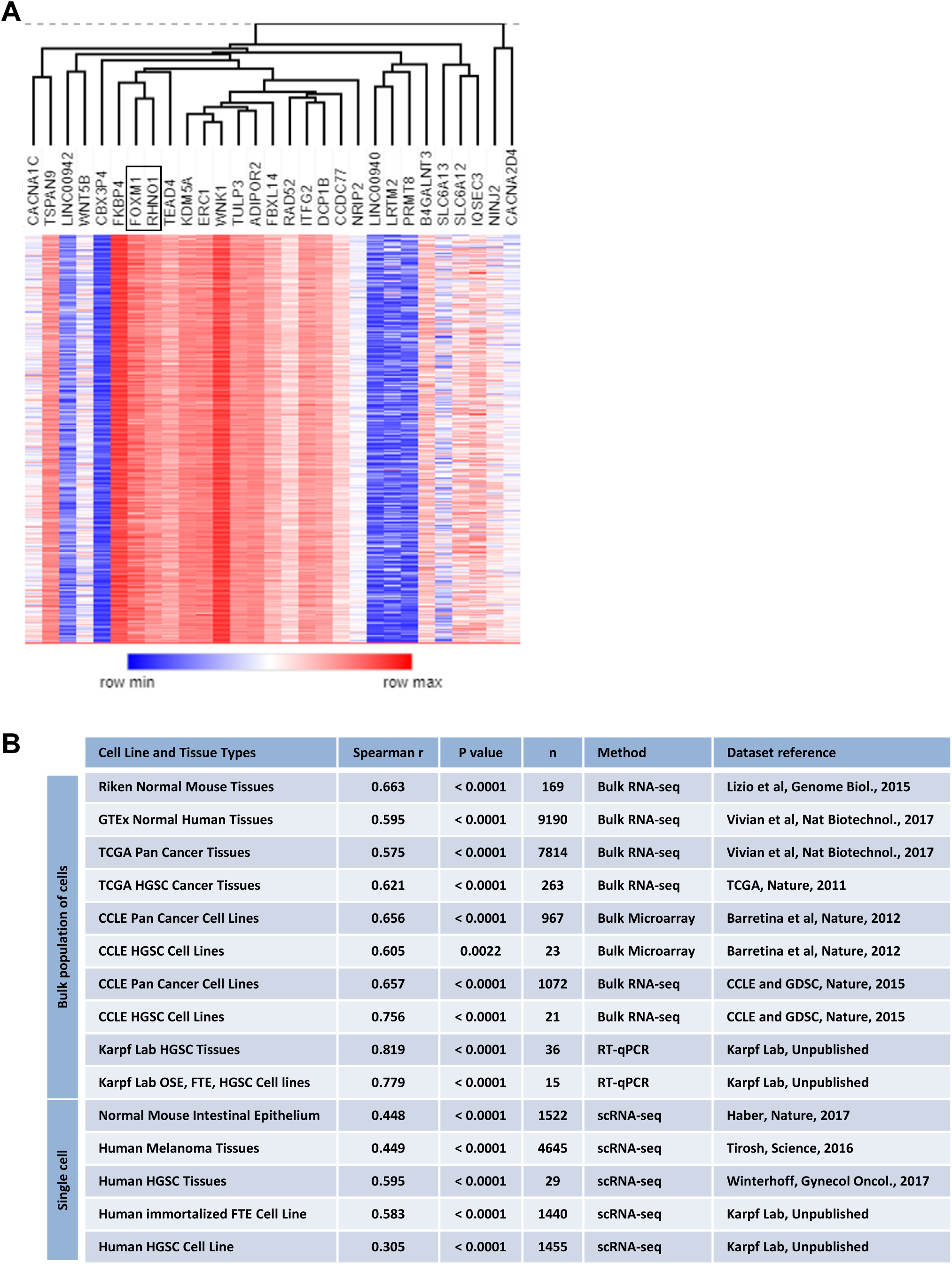
Correlations of 12p13.33 amplicon genes in HGSC, and correlated *FOXM1* and *RHNO1* expression in tissues and cells. **A.** Hierarchical clustering dendogram illustrating **the** correlation of 33 genes in the TCGA HGSC 12p13.33 amplicon. **B.** *FOXM1* and *RHNO1* mRNA correlations in bulk RNA-seq, RT-qPCR and single cell RNA-seq data sets from normal and cancer tissues and cell lines, including human and mouse.

The data presented above suggested that *FOXM1* and *RHNO1* are co-regulated. In agreement, we observed highly significant correlation of *FOXM1* and *RHNO1* expression in normal and cancer tissues, and in both mouse and human samples (**Figure 1B**). Although bidirectional gene partners are often co-expressed (Bornelov et al., 2015; Wakano et al., 2012; Yang et al., 2007); it is unclear if this correlation is maintained at the level of individual cells. To test this for *FOXM1* and *RHNO1*, we analyzed published and newly generated single cell RNA-seq (scRNA-seq) datasets (Haber et al., 2017; Tirosh et al., 2016; Winterhoff et al., 2017). Notably, there was significant correlation between *FOXM1* and *RHNO1* in single cells, and the correlation in bulk and single cell sample populations were similar (**Figure 1B**).

### *FOXM1* and *RHNO1* expression are both elevated in pan-cancer

Although *FOXM1* is widely overexpressed in cancer, *RHNO1* expression in cancer tissues has not been determined (Barger et al., 2019; Kim et al., 2010). We used the TOIL TCGA dataset to compare *FOXM1* and *RHNO1* mRNA expression in normal tissues versus primary, metastatic and recurrent cancer tissues, and found that both *FOXM1* and *RHNO1* are overexpressed in each category of tumor samples **(Figure S4A)**. Although both genes are overexpressed in cancer, we noted that *FOXM1* expression was lower than *RHNO1* in normal tissues and higher in tumor tissues, and the *FOXM1/RHNO1* expression ratio was elevated in tumors **(Figure S4A-B)**. We hypothesized that this might reflect increased proliferation in tumors, as FOXM1 is a proliferation associated transcription factor (Ye et al., 1997). In agreement, after normalization to the canonical cell proliferation marker *MKI67* (Whitfield et al., 2006), the *FOXM1*/*RHNO1* expression ratio was similar in normal and tumor tissues **(Figure S4B)**.

As mentioned above, the F/R-BDP contains a CGI, a potential region for differential DNA methylation, which was previously shown to regulate BDP activity (Shu et al., 2006). We hypothesized that differential DNA methylation might contribute to the *FOXM1* and *RHNO1* expression differences seen in normal and cancer tissues. We performed *in silico* analysis of DNA methylome data from human cancer cell lines (CCLE) and normal and cancer tissues (TCGA), and found that the F/R-BDP CGI was hypomethylated in all datasets (**Figure S5**). As 450K arrays did not encompass all CpG sites in the F/R-BDP, we conducted bisulfite clonal sequencing of normal ovary and ovarian cancer tissues. The F/R-BDP was fully hypomethylated in both tissues (**Figure S6**). These data indicate that altered DNA methylation does not account for the increased *FOXM1* and *RHNO1* mRNA expression observed in human cancer.

### *FOXM1* and *RHNO1* are expressed in primary and recurrent HGSC, and expression correlates with genomic copy number

FOXM1 overexpression and pathway activation are highly prevalent in HGSC (Barger et al., 2019; Barger et al., 2015; Cancer Genome Atlas Research, 2011). FOXM1 is overexpressed in HGSC compared to fallopian tube epithelial (FTE) tissues and cells, and copy number alterations is an important mechanism accounting for this (Barger et al., 2019; Barger et al., 2015). Here we analyzed both *FOXM1* and *RHNO1* expression in HGSC and determined their relationship to genomic copy number. Using TOIL TCGA HGSC and GTEx fallopian tube and ovary RNA-seq data, we found that both *FOXM1* and *RHNO1* are highly increased in primary HGSC (**Figure 2A**). In agreement, *FOXM1* and *RHNO1* expression were significantly elevated in HGSC cells compared to control cells (FTE and OSE) (**Figure 2B**).

**Figure 2.**
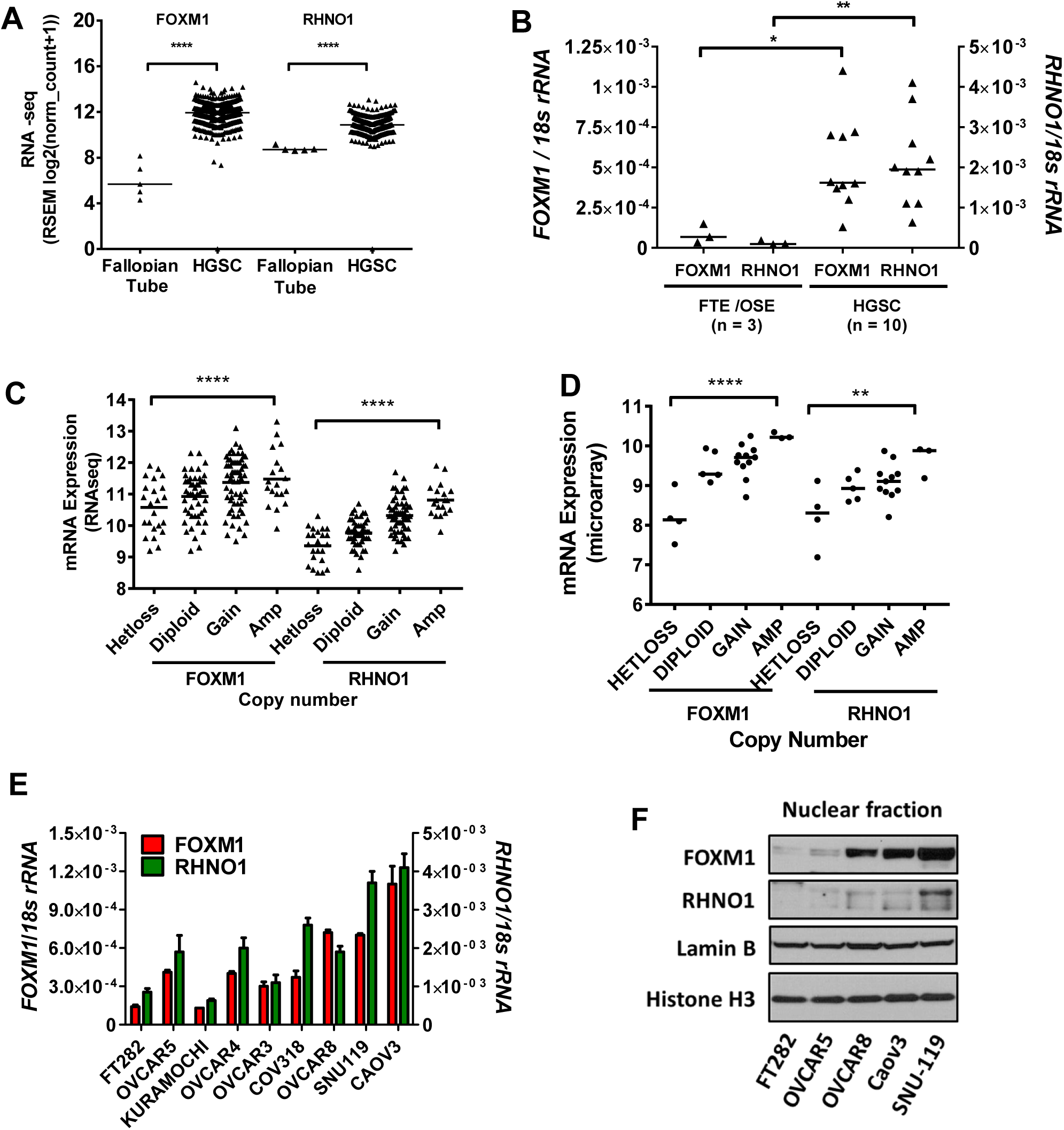
*FOXM1* and *RHNO1* expression and copy number in HGSC tissues and cells. **A.** *FOXM1* and *RHNO1* expression (RNA-seq) in TCGA HGSC tissue as compared to GTEx normal fallopian tube tissues. **B.** *FOXM1* and *RHNO1* mRNA expression from FTE and OSE cell lines compared to HGSC cell lines as measured by RT-qPCR. **C.** *FOXM1* and *RHNO1* mRNA expression (RNA-seq) correlated with copy number (GISTIC) in TCGA HGSC tissues. The p value for ANOVA with post-test for linear trend is shown. Lines represent group medians. **D.** *FOXM1* and *RHNO1* mRNA expression (microarray) correlated with copy number (GISTIC) in CCLE HGSC cell lines. The p value for ANOVA with post-test for linear trend is shown. Lines represent group medians. **E.** *FOXM1* and *RHNO1* mRNA expression were measured RT-qPCR. **F.** FOXM1 and RHNO1 protein expression were measured by Western blot. Lamin B and Histone H3 are shown as loading controls. **E** and **F** plots cell types with increasing *FOXM1* and *RHNO1* copy number from left to right. T-test *P* value is shown. *P* value designation: **** < 0.0001, *** < 0.001, ** < 0.01, * < 0.05.

As disease recurrence is a major clinical issue in ovarian cancer, we analyzed *FOXM1* and *RHNO1* expression in primary vs. recurrent chemoresistant HGSC (Kreuzinger et al., 2017; Patch et al., 2015). There was strong concordance between *FOXM1* and *RHNO1* expression in both primary and recurrent samples (**Figure S7**). In addition, the *FOXM1/RHNO1* expression ratio increased in recurrent HGSC and, in the Australian data set (Patch et al., 2015), 5/11 patients had increased *FOXM1* and *RHNO1* expression at recurrence (**Figures S8-9**). Thus, *FOXM1* and *RHNO1* maintain balanced expression in both primary and recurrent HGSC, and are frequently coordinately upregulated in recurrent chemoresistant HGSC.

TCGA HGSC and CCLE HGSC cell line data revealed a progressive increase in *FOXM1* and *RHNO1* expression with genomic copy number **(Figures 2C-D)**. Additionally, FOXM1 and RHNO1 mRNA and protein expression in immortalized FTE cells and HGSC cell lines increased with genomic copy number **(Figures 2E-F).**

### The F/R-BDP promotes balanced *FOXM1* and *RHNO1* gene expression

To determine whether *FOXM1* and *RHNO1* expression are regulated by their putative BDP, we first determined the transcriptional start sites (TSS) for both genes, using immortalized FTE cells and two HGSC cell lines. The experimentally defined TSS and intergenic distance were similar to those predicted by NCBI, and indicated an BDP of approximately 200 bp, well within the generally defined distance of < 1kb assigned to BDPs (Trinklein et al., 2004) (**Figure S10**). We cloned the putative F/R-BDP into a reporter construct where the *FOXM1* promoter drives Renilla luciferase and the *RHNO1* promoter drives Firefly luciferase (**Figure 3A**). Transfection into immortalized FTE cells and a panel of HGSC cell lines revealed a direct correlation between *FOXM1* and *RHNO1* promoter activity (**Figure 3B**). Additionally, F/R-BDP activity of the cognate promoter directly correlated with endogenous *FOXM1* and *RHNO1* mRNA expression (**Figure 3C-D)**. To assess F/R-BDP activity at a single cell level, we cloned the F/R-BDP into a reporter construct where the *FOXM1* promoter drives green fluorescent protein (GFP) expression and the *RHNO1* promoter drives red fluorescent protein (RFP) expression **(Figure S11A)**. In agreement with luciferase assay data, there was a significant direct correlation between GFP and RFP expression in 293 cells transfected with the F/R-BDP construct **(Figure S11B-E)**.

**Figure 3.**
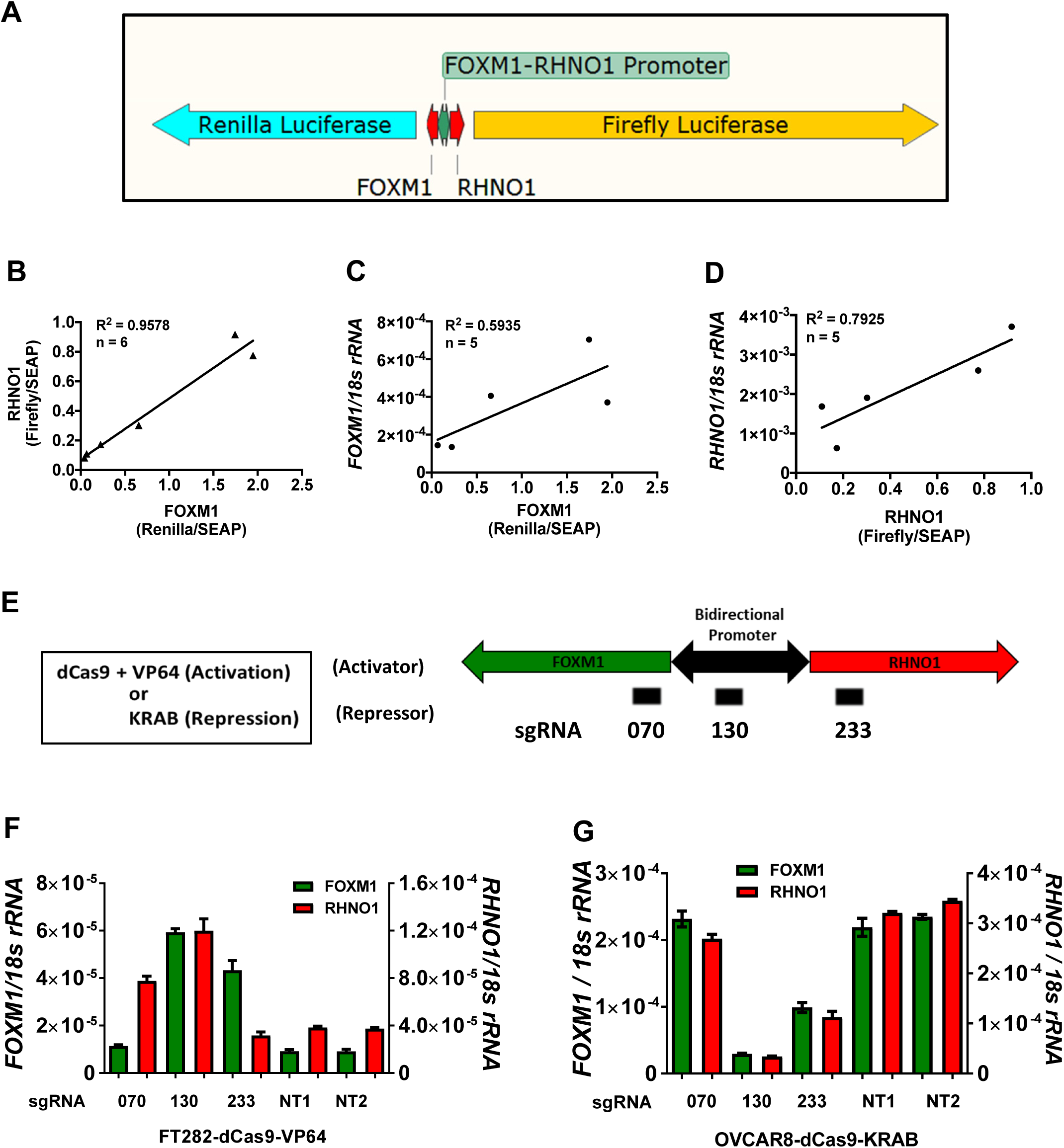
*FOXM1* and *RHNO1* bidirectional promoter (F/R-BDP) regulation. **A-D.** F/R-BDP reporter assay. **A.** F/R-BDP reporter construct with the promoter colored in green. *FOXM1* drives renilla luciferase and *RHNO1* drives firefly luciferase. Secreted embryonic alkaline phosphatase (SEAP) was co-transfected and used as an internal normalization control. **B.** Correlation of *FOXM1* and *RHNO1* promoter activity in a panel of FTE and HGSC cells. **C.** Correlation of *FOXM1* promoter activity with endogenous *FOXM1* mRNA measured by RT-qPCR. **D.** Correlation of *RHNO1* promoter activity with endogenous *RHNO1* mRNA measured by RT-qPCR. **E-G.** CRISPR mediated activation and repression of the endogenous F/R-BDP. **E.** F/R-BDP reporter in blue and guide RNAs indicated below. **F.** CRISPR mediated activation of the F/R-BDP in FTE cells expressing synergistic activation mediator (SAM) and a guide RNA targeting the BDP region and the corresponding changes in mRNA expression as measured by RT-qPCR. **G.** CRISPR mediated inhibition of the F/R-BDP in HGSC cells expressing Krüppel associated box (KRAB) transcriptional repressor and a guide RNA targeting the bidirectional promoter and the corresponding changes in mRNA expression as measured by RT-qPCR. NT1 and NT2 – non-targeting control guide RNAs.

We next utilized a CRISPR activation system to study the endogenous F/R-BDP. This system uses a single guide RNA (sgRNA) to recruit a catalytically dead Cas9 (dCas9) fused to the VP64 transcriptional activator (Joung et al., 2017). We designed three sgRNA to target the F/R-BDP or flanking regions (**Figure 3E**), and introduced the system into FT282 cells, which have low endogenous *FOXM1* and *RHNO1* expression. Interestingly, sgRNA targeting outside the F/R-BDP, upstream of the TSS for one gene and within the gene body of the other gene (i.e. 070 or 233), only induced the expression of the distal gene (**Figure 3F**). In contrast, the sgRNA directly targeting the F/R-BDP (i.e. 130) induced the expression of both *FOXM1* and *RHNO1* (**Figure 3F**). In a converse experiment, we used a CRISPR-KRAB repressor (i.e. CRISPR inhibition or CRISPRi system) (Thakore et al., 2015) to target the F/R-BDP or flanking regions in an HGSC cell line with high endogenous *FOXM1* and *RHNO1* expression (OVCAR8). Notably, the guide RNA targeting the BDP (i.e. 130) efficiently repressed both *FOXM1* and *RHNO1* (**Figure 3G**). Together, these data reveal that the F/R-BDP promotes balanced *FOXM1* and *RHNO1* expression.

### FOXM1 and RHNO1 cooperatively promote HGSC cell growth and survival and regulate cell cycle progression

Based on the data presented above, we hypothesized that FOXM1 and RHNO1 may have cooperative functions, particularly in cancer cells. More specifically, we hypothesized that they may cooperatively promote cell growth and survival, as FOXM1 is known to promote cell cycle progression and DNA repair, while RHNO1 promotes cell survival, the DNA replication stress response, and DDR (Barger et al., 2015; Cotta-Ramusino et al., 2011; Kim et al., 2010; Lindsey-Boltz et al., 2015; Zona et al., 2014). To test these ideas, we engineered OVCAR8 and CAOV3 HGSC cells for inducible FOXM1 and RHNO1 shRNA knockdown and measured clonogenic survival. FOXM1 or RHNO1 knockdown each reduced the clonogenic growth of both cell types (**Figure 4**). To validate these observations, we used CRISPR-Cas9 to knockout FOXM1 or RHNO1 in both cell types, and we again observed a significant reduction of HGSC clonogenic growth **(Figure S12A-D)**. CRISPR knockout of FOXM1 or RHNO1 also increased apoptosis, supporting a role for these proteins in cell survival **(Figure S12E)**. To determine the combined effects of FOXM1 and RHNO1 depletion, we engineered OVCAR8 cells for dual FOXM1 and RHNO1 knockdown (**Figure 5A**). Dual knockdown reduced clonogenic growth to a greater degree than either single knockdown, suggesting functional cooperativity (**Figure 5B**).

**Figure 4.**
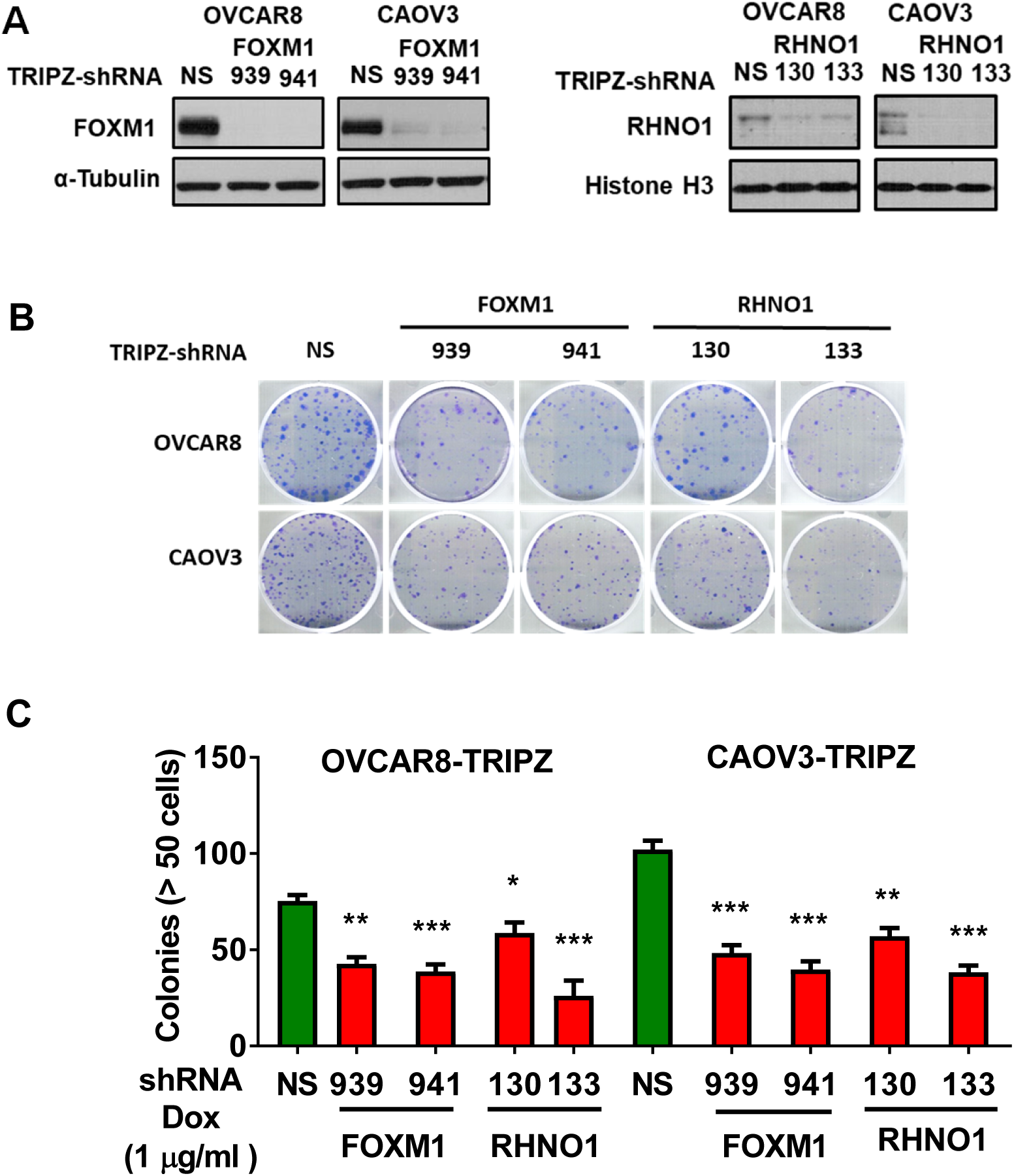
Clonogenic survival of HGSC cells following FOXM1 or RHNO1 knockdown. OVCAR8 and CAOV3 cells engineered for dox-inducible FOXM1 or RHNO1 shRNA knockdown were seeded for protein extractions or clonogenic survival assays. **A.** Cells were grown in the presence of dox for 72 hours and protein was harvested for Western blot analysis to confirm knockdown efficiency. **B.** Cells were seeded into a 6-well dish, in triplicate, at a density of 500 or 1000 cells, respectively. Dox was added at the time of seeding and media containing dox was replenished every 48 hours. Clonogenic survival was measured at 12 and 14 days, respectively, after the cells were fixed with methanol and stained with crystal violet. **C.** Colonies containing more than 50 cells were counted and clonogenic survival was quantified as an average of the replicates. NS = non-targeting shRNA. t test *P* value is shown. *P* value designation: **** < 0.0001, *** < 0.001, ** < 0.01, * < 0.05.

**Figure 5.**
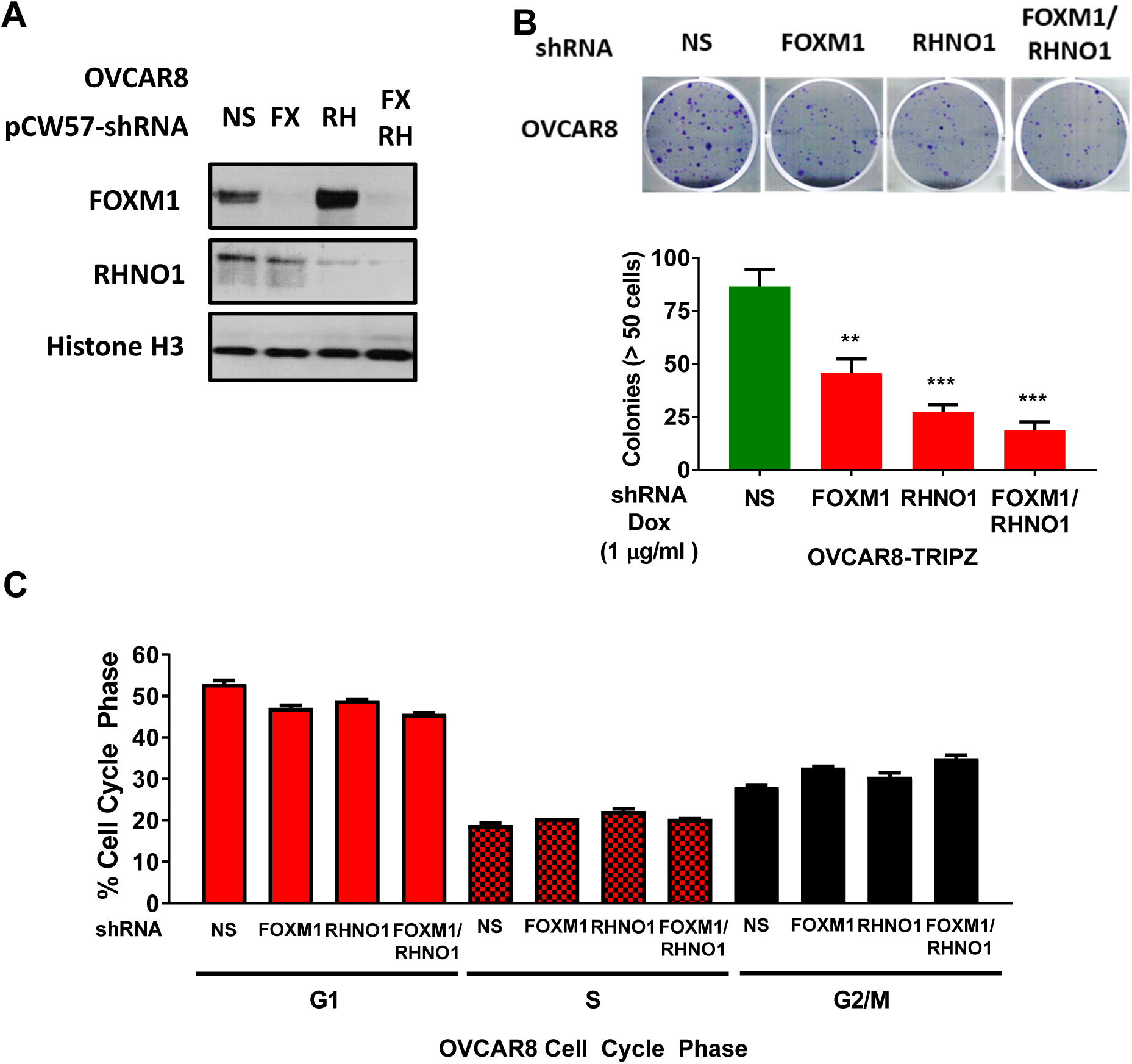
Clonogenic survival and cell cycle of HGSC cells following combined FOXM1 and RHNO1 knockdown. OVCAR8 cells engineered for dox-inducible FOXM1 and/or RHNO1 shRNA knockdown were seeded for protein or clonogenic survival. **A.** Cells were grown in the presence of dox for 72 hours to harvest protein followed by Western blot analysis to confirm knockdown efficiency. **B.** Cells were seeded into a 6-well dish, in triplicate, at a density of 500 cells. Dox was added at the time of seeding and media containing dox was replenished every 48 hours. Clonogenic survival was measured at 12 days, cells were fixed with methanol and stained with crystal violet. **C.** Colonies containing more than 50 cells were counted and clonogenic survival was quantified as an average of the replicates. NS = non-targeting shRNA. t test *P* value is shown. *P* value designation: **** < 0.0001, *** < 0.001, ** < 0.01, * < 0.05.

### FOXM1 regulates cell cycle and the G2/M checkpoint in HGSC cells

FOXM1 mediates cellular phenotypes through its function as a transcription factor. To understand how loss of FOXM1 reduced HGSC cell growth, we performed RNA-seq analysis on OVCAR8 and CAOV3 cells following FOXM1 knockdown (**Figure S13A-B**). Gene set enrichment analyses (GSEA) revealed significant enrichment of cell cycle gene expression signatures in both cell types and, in particular, enriched mitotic and G2/M checkpoint signatures (**Figure S13C-H**). FOXM1 knockdown did not show enriched DNA repair expression signatures, as reported in other cell types, suggesting cell context-dependent FOXM1 transcriptional programs (Fang et al., 2018; Tan et al., 2007). We validated target gene expression using RT-qPCR and found that *CCNB1* was downregulated in both CAOV3 and OVCAR8 cells after FOXM1 knockdown (**Figure S13I-J**). However, FOXM1 knockdown did not alter expression of the previously reported target DNA repair gene *BRCA2* (Kwok et al., 2010) (**Fig S13K**). These data suggest that, in HGSC cells, FOXM1 primarily regulates genes involved in cell cycle progression.

To validate the functional impact of FOXM1 on HGSC cell cycle, and to define RHNO1’s impact, we determined the cell cycle profiles of OVCAR8 cells sustaining FOXM1 and/or RHNO1 knockdown. FOXM1 knockdown caused an increased proportion of G2/M cells (**Figure 5C**). In contrast, RHNO1 knockdown led to an increased G1/S proportion, consistent with reduced S phase progression, in agreement with previous work (Cotta-Ramusino et al., 2011; Kim et al., 2010) (**Figure 5C**). Surprisingly, dual FOXM1 and RHNO1 knockdown showed a greater increase in G2/M proportion as compared to FOXM1 knockdown alone (**Figure 5C**).

### RHNO1 interacts with 9-1-1 and promotes ATR-CHK1 signaling, HR, and cell survival in HGSC cells

RHNO1 interacts with the 9-1-1 complex and TOPBP1 to promote ATR activation (Cotta-Ramusino et al., 2011; Lindsey-Boltz et al., 2015). This function has not been assessed in HGSC cells, or after treatments that specifically cause DNA replication stress. To address this, we initially overexpressed HA-tagged RHNO1-WT and RHNO1-SWV mutant in 293T cells and performed Co-IP experiments. RHNO1-SWV has mutations in the APSES DNA binding domain that disrupt its interaction with 9-1-1 but not TOPBP1 (**Figure S14A**) (Cotta-Ramusino et al., 2011). Consistent with the prior report, we observed that RHNO1-WT interacted with RAD9 and RAD1 while RHNO1-SWV was deficient for these interactions (**Figure S14B**). Next, we investigated RHNO1 localization and interactions in HGSC cells. In OVCAR8 cells, RHNO1 was present at chromatin at baseline, while treatment with hydroxyurea (HU), a classical inducer of DRS, further increased its chromatin localization (**Figure S14C**). To test RHNO1 interaction with 9-1-1 in OVCAR8 cells, we depleted endogenous RHNO1 using shRNA and reconstituted cells with shRNA-resistant HA-tagged RHNO1-WT and RHNO1-SWV, in a dox-inducible manner. The data indicated that RHNO1-WT interacts with 9-1-1 and TOPBP1, but RHNO1-SWV only interacts with TOPBP1 (**Figure S14D**). To address the function of RHNO1 in ATR-CHK1 signaling in OVCAR8, we depleted RHNO1 using dox-inducible shRNA and measured P-CHK1-S345 (Niida et al., 2007). We observed a ∼2-fold reduction of P-CHK1-S345 in RHNO1 depleted cells, similar to that reported previously for RHNO1 (Cotta-Ramusino et al., 2011; Lindsey-Boltz et al., 2015) (**Figure S14E**). By impairing ATR-CHK1 signaling, RHNO1 knockdown may increase DNA damage. To test this, we measured γ-H2AX in different cell cycle phases, and observed a significant increase of γ-H2AX- in both S and G2 phases with RHNO1 knockdown (**Figure S14F**). Moreover, COMET analyses revealed significantly increased DNA strand breaks in RHNO1 knockdown cells (**Figure S14G**). These data establish that RHNO1 functions in the replication stress response and DNA damage protection in HGSC cells.

Next, to test whether 9-1-1 interaction is required for the cell survival function of RHNO1, we knocked down endogenous RHNO1 with dox-inducible shRNA and simultaneously expressed dox-inducible, shRNA-resistant, RHNO1-WT or RHNO1-SWV, and measured OVCAR8 clonogenic survival (**Figure S15A-B**). Importantly, only RHNO1-WT partially rescued the defect in clonogenic survival (**Figure S15C-D**). To determine if reduced survival following RHNO1 loss is retained in non-cancer cells, we used CRISPR to knock out RHNO1 in immortalized FTE cells (**Figure S15E-F**). Importantly, RHNO1 knockout did not significantly decrease FTE cell growth and viability (**Figure S15G**). These data suggest that cancer cells may have an increased dependency on RHNO1 for growth and survival compared to non-transformed cells, and demonstrate that 9-1-1 interaction is critical for RHNO1’s role in cell survival.

### FOXM1 and RHNO1 cooperatively promote HR

FOXM1 is reported to promote HR (Khongkow et al., 2014; Maachani et al., 2016; Monteiro et al., 2013; Park et al., 2012); however, FOXM1 depletion in OVCAR8 cells did not appear to alter the expression of DNA damage response genes, including *BRCA2* (**Figure S13**). By virtue of its role in the DRS response, RHNO1 promotes HR (Cotta-Ramusino et al., 2011). Thus, based on their genomic arrangement, co-regulation, and reported functions, we hypothesized that FOXM1 and RHNO1 may cooperatively promote HR. To test this, we first used DR-GFP assays in U2OS cells (Gunn et al., 2011; Gunn and Stark, 2012). We engineered these cells for inducible FOXM1 and RHNO1 knockdown alone and in combination, or for RAD51 knockdown as a positive control for HR impairment (**Figure 6A-B**) (Cotta-Ramusino et al., 2011). Notably, RHNO1 knockdown decreased HR efficiency but FOXM1 knockdown did not. However, unexpectedly, dual knockdown showed greater impairment in HR compared to RHNO1 knockdown alone, suggesting functional cooperativity (**Figure 6E**). We next used OVCAR8 DR-GFP cells to validate these observations in HGSC cells. OVCAR8 cells have a methylated *BRCA1* promoter but remain HR proficient (Huntoon et al., 2013). Similar to U2OS cells, FOXM1 loss slightly reduced HR and RHNO1 loss substantially reduced HR efficiency, while dual loss resulted in the most signification reduction (**Figure 6C,D,F**).

**Figure 6.**
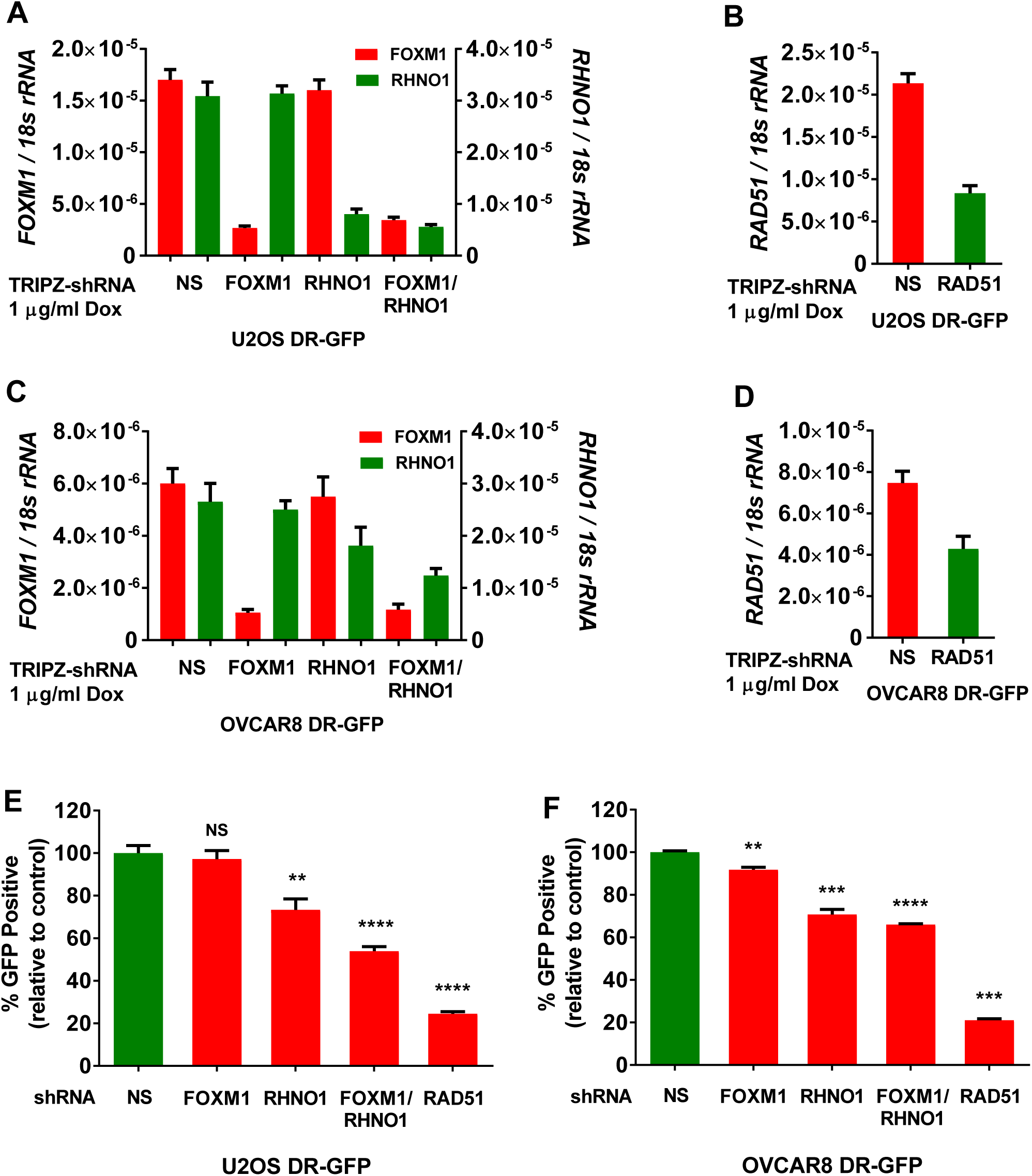
FOXM1 and RHNO1 promote HR in U2OS and OVCAR8 cells. Engineered DR-GFP cell lines were treated with doxycycline to induce shRNA expression and gene knockdown was measured by RT-qPCR. **A,C**. *FOXM1* and *RHNO1* mRNA expression. **B,D.** *RAD51* mRNA expression. **E-F**. HR repair rate of I-SceI-induced DSBs was measured in **E.** U2OS and **F.** OVCAR8 DR-GFP cells. Each value is expressed relative to the percentage of GFP-positive cells in I-SceI-transfected (U2OS) and transduced (OVCAR8) control cells. Student’s *t*-test was used for comparisons and RAD51 knockdown was used as a positive control for HR impairment. All results are shown as mean ± SE from three independent replicates (\**P* < 0.05, \*\**P* < 0.01, \*\*\**P* < 0.005).

### FOXM1 and RHNO1 depletion sensitizes HGSC cells to the PARPi olaparib and mitigates acquired olaparib resistance

HR is impaired in approximately half of primary HGSC, which confers PARPi sensitivity (Cancer Genome Atlas Research, 2011; Farmer et al., 2005; Lord et al., 2015). Based on the data presented above, we hypothesized that loss of FOXM1 and/or RHNO1 may potentiate the response to PARPi, particularly in HR proficient HGSC cells. To test this, we first utilized OVCAR8 cells with dox-inducible single and dual knockdown of FOXM1 and RHNO1 and determined the IC_50_ of the PARPi olaparib, using AlamarBlue assays. While each single knockdown moderately sensitized OVCAR8 to olaparib, dual knockdown led to the greatest increase in sensitivity (**Figure 7A-B**). Consistently, there was increased DNA damage in dual knockdown cells treated with olaparib (**Figure 7C**). Furthermore, dual FOXM1 and RHNO1 knockdown resulted in an increased proportion of cells in G2/M with olaparib treatment compared to either single knockdown (**Figure 7D**). To directly test whether F/R-BDP activity controls olaparib sensitivity, we used a CRISPRi strategy in OVCAR8 cells. F/R-BDP repression sensitized OVCAR8 to olaparib to a similar extent seen with dual FOXM1 + RHNO1 shRNA knockdown (**Figure 8A-C**). In addition, F/R-BDP repression sensitized OVCAR8 cells to olaparib-mediated apoptosis (**Figure 8D**) and increased G2/M arrest (**Figure 8E**).

**Figure 7.**
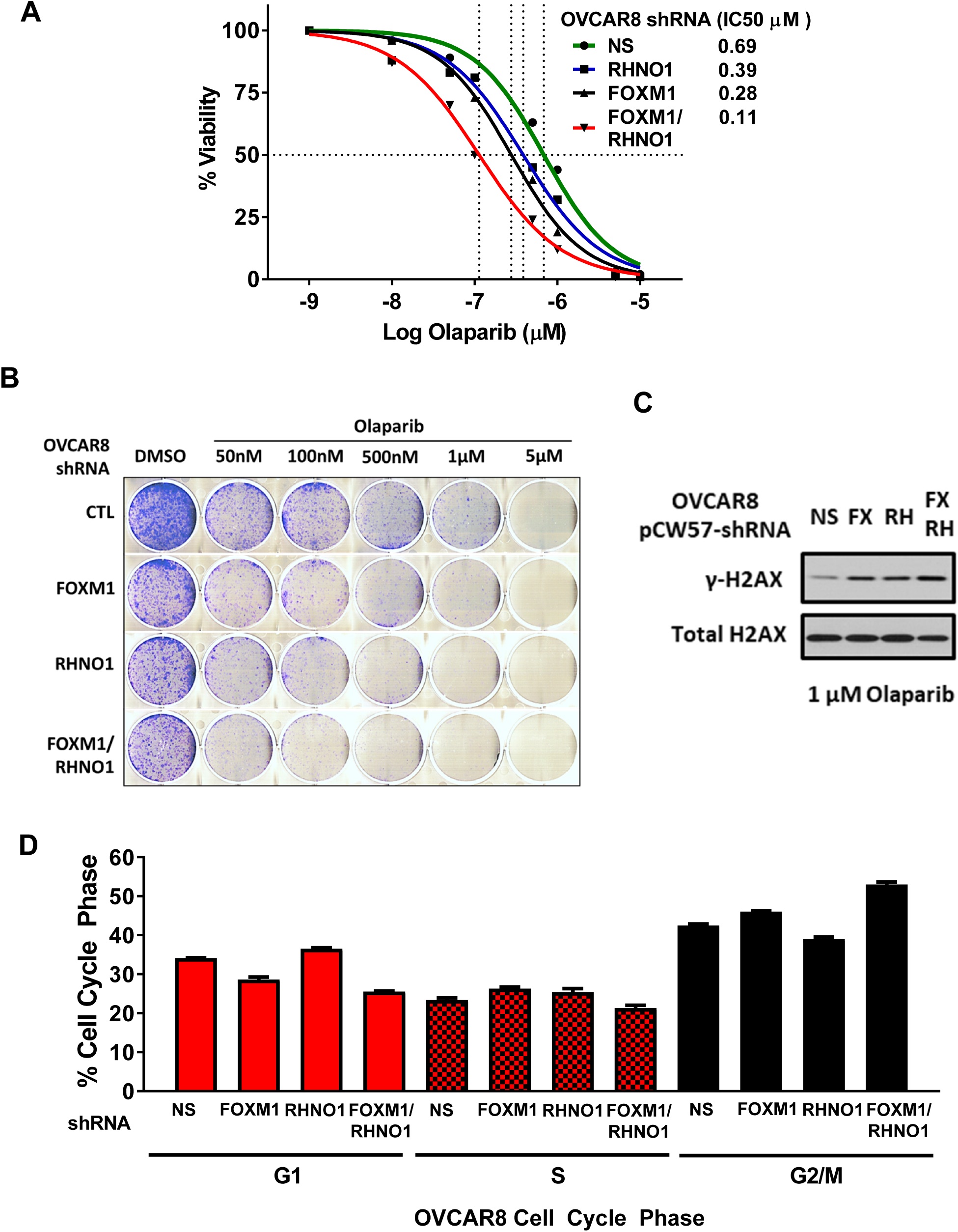
FOXM1 and RHNO1 knockdown sensitizes OVCAR8 cells to olaparib. OVCAR8 cells engineered for dox-inducible FOXM1 and/or RHNO1 shRNA knockdown were grown in the presence of dox for 72 hours prior to seeding cells for olaparib IC50 or clonogenic survival. **A.** Cells were harvested then seeded in 96-well plates in quadruplicate in the presence of dox at a density of 500 cells per well. Twenty-four hours later cells received media containing dox and vehicle or olaparib, and this was repeated every 48 hours. Cell viability was measured at 8 days using AlamarBlue and the IC50 for olaparib was determined. **B**. Cells were harvested and seeded into a 6-well dish, in triplicate, in the presence of dox at a density of 5000 cells per well. Twenty-four hours later cells received media containing dox and vehicle or olaparib, and this was repeated every 48 hours. Media containing dox was replenished every 48 hours. After 8 days of growth, cells were fixed with methanol and stained with crystal violet. NS = non-targeting shRNA. **C.** The indicated protein extracts were used for gH2AX western blot. **D.** The indicated samples were used for cell cycle analyses using PI staining followed by flow cytometry analyses.

**Figure 8.**
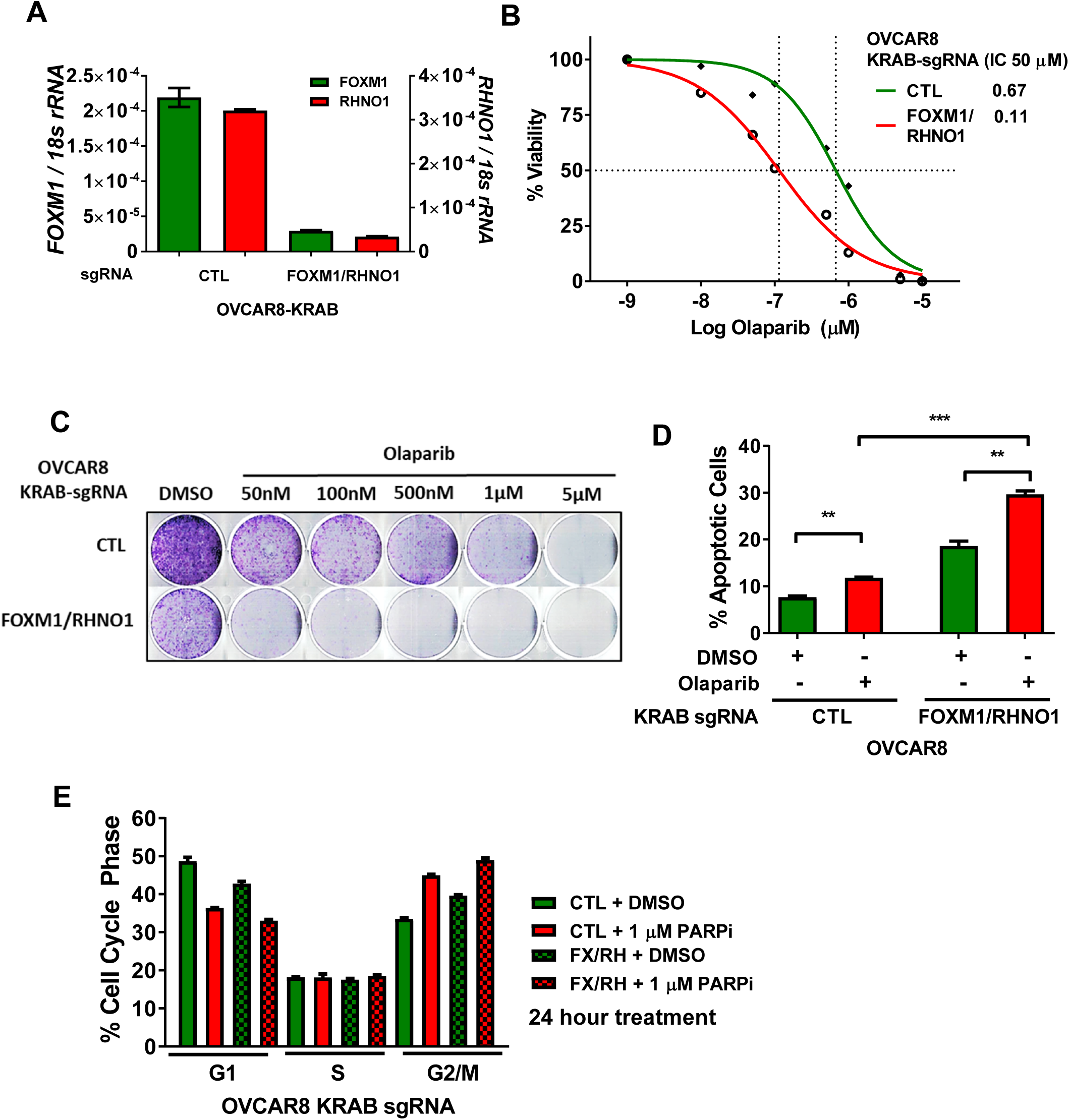
CRISPR interference of the F/R-BDP sensitizes OVCAR8 cells to olaparib. **A.** OVCAR8 cells expressing KRAB transcriptional repressor and a guide RNA targeting the F/R-BDP, and the corresponding changes in mRNA expression as measured by RT-qPCR. NT1 – non-targeting guide RNA. **B.** Cells were seeded in 96-well plates in quadruplicate at a density of 500 cells per well. Twenty-four hours later cells received media containing vehicle or olaparib, and this was repeated every 48 hours. Cell viability was measured at 8 days using AlamarBlue and the IC50 for olaparib was determined**. C**. Cells were seeded into a 6-well dish, in triplicate, at a density of 5,000 cells per well. Twenty-four hours later cells received media containing vehicle or olaparib, and this was repeated every 48 hours. After 8 days of growth, cells were fixed with methanol and stained with crystal violet. **D.** Apoptosis were measured by Annexin V staining and FACS. **E.** Cell cycle was measured with PI staining and flow cytometry.

Although HR-deficient tumor cells are selectively sensitive to PARPi, these tumors often develop PARPi resistance (Bitler et al., 2017; Fojo and Bates, 2013; Lord et al., 2015; Sonnenblick et al., 2015). To examine the potential function of the F/R-BDP in acquired PARPi resistance, we used isogenic derivatives of UWB1.289 (UWB1), a BRCA1-null HGSC cell line with *in vitro* acquired olaparib resistance (Yazinski et al., 2017). Two PARPi-resistant UWB1 derivatives, SyR12 and SyR13, are dependent on ATR for cell survival in the presence of PARPi, and GSEA analysis showed enrichment for G2/M checkpoints and ATR signaling in these cells (Yazinski et al., 2017). Notably, CRISPRi targeting the F/R-BDP silenced *FOXM1* and *RHNO1* mRNA expression, and re-sensitized SyR12 and SyR13 to olaparib (**Figure 9**). F/R-BDP repression did not completely restore olaparib sensitivity, similar to the effects seen with ATRi (Yazinski et al., 2017).

**Figure 9.**
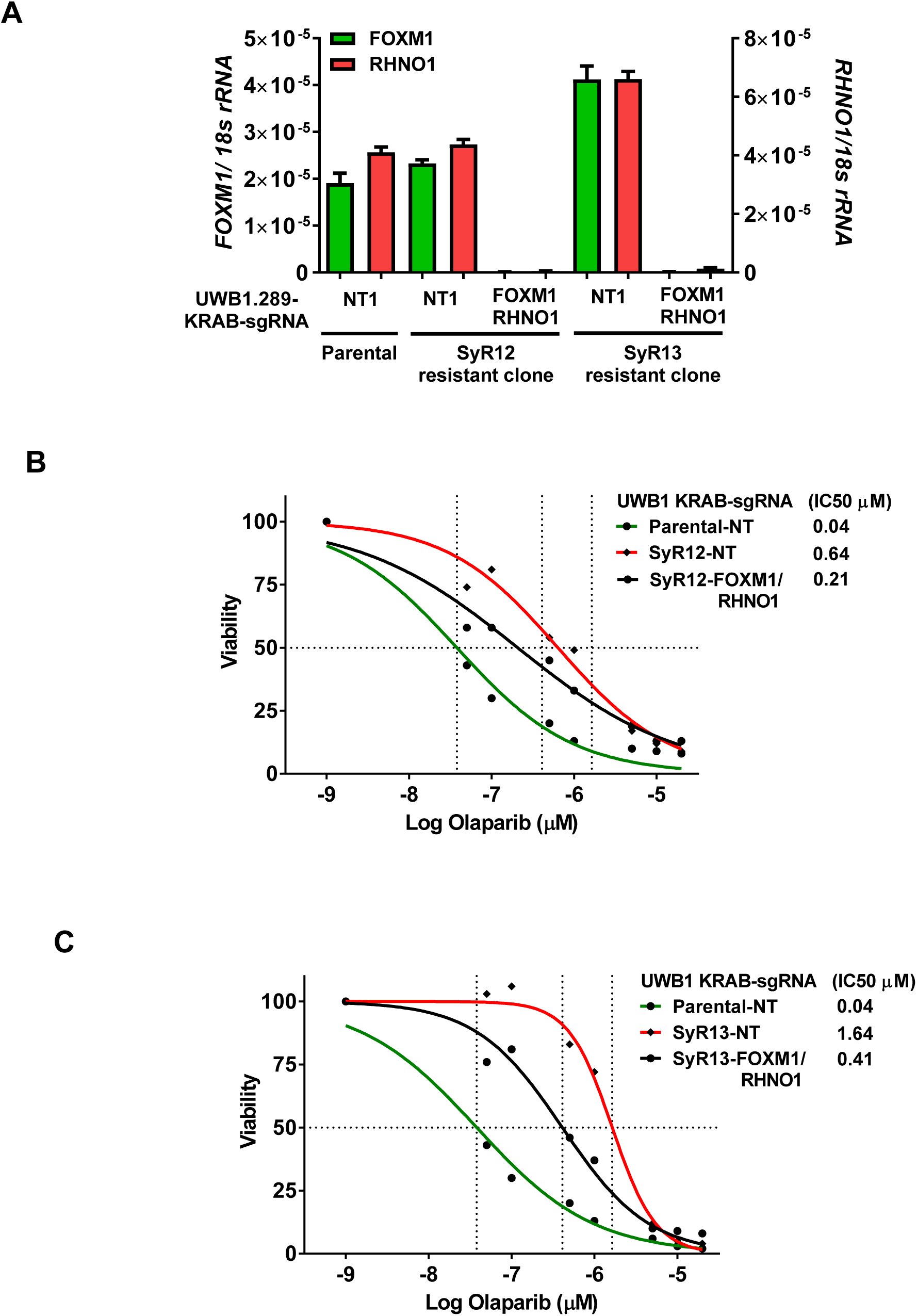
CRISPR interference of the F/R-BDP in UWB1 PARPi-resistant clones restores PARPi sensitivity. UWB1 parental and PARPi resistant cells were engineered to express KRAB transcriptional repressor and a control guide RNA or guide RNA targeting the F/R-BDP. **A.** Cells were seeded and grown for 72 hours to harvest RNA followed by RT-qPCR to confirm knockdown efficiency**. B-C.** UWB1 parental KRAB and (**B)** SyR12 KRAB or (**C)** SyR13 KRAB cells were seeded in 96-well plates in quadruplicate at a density of 750 cells per well. Twenty-four hours later cells received media containing vehicle or olaparib, and this was repeated every 48 hours. Cell viability was measured at 8 days using AlamarBlue and the IC50 for olaparib was determined. NT1 – non-targeting guide RNA.

## Discussion

Analysis of the 12p13.33 HGSC amplicon revealed that *FOXM1* is arranged head-to-head with *RHNO1*, a recently characterized gene whose encoded protein promotes ATR-CHK1 signaling through interaction with the 9-1-1 complex (Cotta-Ramusino et al., 2011; Lindsey-Boltz et al., 2015). Genomic data revealed strong correlation between *FOXM1* and *RHNO1* mRNA expression in normal, HGSC and pan-cancer tissues, and this was confirmed in bulk RNA-seq data from our independent HGSC tumor bank and HGSC cell lines, and in scRNA-seq data of immortalized FTE cells and HGSC cells. We demonstrate that the *FOXM1* and *RHNO1* intergenic space functions as a BDP both in immortalized FTE and HGSC cells. Functionally, depletion of FOXM1 and RHNO1 reduced HR efficiency and clonogenic growth and survival in HGSC cells. Furthermore, in HGSC cells FOXM1 promoted cell cycle progression while RHNO1 binds 9-1-1 and TOPBP1 and promotes ATR-CHK1 signaling, HR, and cell survival. Perhaps most importantly, FOXM1 and RHNO1 cooperatively promote PARPi resistance, both in naïve HR-competent HGSC cells and in HR-deficient HGSC cells with acquired PARPi resistance. These data support additional studies of FOXM1 and RHNO1 to further the understanding of their possible cooperative role in HGSC, and to determine if they mediate cooperative functions in normal cells and other cancers. More broadly, our findings support the study of other bidirectional gene pairs to investigate their potential cooperative functional contribution to cancer. Recent data have shown that genetic and epigenetic alterations of bidirectional genes, including copy number alterations, are selected against in cancer (Thompson et al., 2018). Thus, bidirectional gene pairs that show coincident alterations in cancer might have increased functional relevance to cancer.

HGSC tissues and cells show robust overexpression of both *FOXM1* and *RHNO1*. Genomic amplifications and copy number gains promote *FOXM1* and *RHNO1* expression, while DNA methylation does not appear to regulate their expression. It is unknown which transcription factors specifically regulate F/R-BDP activity. Bidirectional promoters are overrepresented with several transcription factor motifs including: GABPA, MYC, E2F1, E2F4, NRF-1, CCAAT, and YY1 (Lin et al., 2007). E2F1 is a strong candidate for F/R-BDP activation, based on our prior data showing that E2F1 activates *FOXM1* expression (Barger et al., 2019; Barger et al., 2015). In agreement, the F/R-BDP contains two E2F motifs.

Surprisingly, while FOXM1 depletion sensitized HGSC cells to olaparib, RNA-seq did not reveal DNA repair genes as transcriptional targets of FOXM1. However, genes involved in the G2/M checkpoint were impacted, and cell cycle analyses showed that FOXM1 knockdown cells accumulated in G2/M, consistent with the role of FOXM1 in G2/M progression (Chen et al., 2013). FOXM1 also plays an important role in mitotic maintenance, and we observed enriched mitotic expression with FOXM1 depletion. In recent studies, decreased levels of FOXM1 was shown to cause mitotic decline, genomic instability, and senescence during human aging (Macedo et al., 2018). Conversely, FOXM1 overexpression was found to promote mitotic defects and genomic instability (Pfister et al., 2018). Together, these data suggest that tight regulation of FOXM1 expression and function are important for maintaining proper mitotic function and genomic stability.

Importantly, olaparib sensitization was greatest in the context of dual FOXM1/RHNO1 knockdown as compared to single gene knockdown. In addition, G2/M accumulation was greatest with dual FOXM1/RHNO1 knockdown, in both the absence and the presence of olaparib. One potential explanation for these findings is that that RHNO1 restricts FOXM1-mediated S to G2 progression, a newly characterized cell cycle checkpoint (Saldivar et al., 2018). Intriguingly, ATR was reported to restrict this checkpoint (Saldivar et al., 2018), suggesting a possible direct role for RHNO1 in regulating FOXM1 function. Additionally, the fact that dual FOXM1/RHNO1 targeting sensitizes cells to olaparib might be explained by the recent observation that PARPi causes increased replication speed, DNA damage, and mitotic defects (Maya-Mendoza et al., 2018). Thus, loss of RHNO1 leads to a reduced efficiency replication stress response, while loss of FOXM1 leads to mitotic defects that might accentuate the DNA damaging effects of olaparib.

Dual FOXM1 and RHNO1 targeting might sensitize HGSC cells to other chemotherapy drugs besides PARPi. This hypothesis is supported by our observation that *FOXM1* and *RHNO1* expression is coordinately upregulated in a sizable proportion of patients with chemoresistant recurrent HGSC (the front line therapy for these patients is carboplatin + paclitaxel). FOXM1 is reported to promote resistance to carboplatin and taxol, and we speculate that RHNO1, which promotes the DDR, might have an additional impact on this phenotype (Carr et al., 2010; Kwok et al., 2010; Nestal de Moraes et al., 2015; Tassi et al., 2017; Wang et al., 2013). In addition, ATR and CHK1 kinases have been shown to have cancer-specific synthetic lethality, suggesting that RHNO1 loss might synergize with a CHK1 inhibitor (Sanjiv et al., 2016). Although there are no reported inhibitors of RHNO1, in theory, ATR inhibitors could be used to target RHNO1 function, given RHNO1’s role in ATR activation (Cotta-Ramusino et al., 2011; Lindsey-Boltz et al., 2015). FOXM1 inhibitors are reported and have been used in preclinical studies (Gartel, 2008; Gormally et al., 2014). Our data showing FOXM1 and RHNO1 co-regulation and functional cooperativity suggests combined targeting of these two proteins as a cancer treatment strategy, possibly using ATRi in combination with FOXM1i. In light of our olaparib data, this combination could be tested with PARPi.

In summary, we demonstrate that the well characterized FOXM1 oncogene is co-expressed with its bidirectional gene partner, RHNO1, and that together the two proteins promote DNA repair, cell survival, and PARPi chemoresistance, to a greater extent than seen with either protein alone. Future studies should seek to further understand their joint contribution to cancer development and progression, and should explore the potential for combinatorial targeting of FOXM1 and RHNO1 as a therapeutic approach.

## Methods

### Human Tissue Samples

Bulk normal ovary (NO) tissues and EOC samples were obtained from patients undergoing surgical resection at Roswell Park Comprehensive Cancer Center (RPCCC) under Institutional Review Board-approved protocols, as described previously (Akers et al., 2014). Pathology specimens were reviewed at RPCCC, and tumors were classified according to World Health Organization criteria (Serov et al., 1973) Flash-frozen bulk tumor tissue samples were crushed using liquid nitrogen pre-chilled mortar and pestles. Total RNA was extracted by TRIzol reagent (Invitrogen) from approximately 20 mg of powdered tissue. The tissue was immediately homogenized in TRIzol by an electric homogenizer with disposable microtube pestles.

### Cell Lines

COV362 and COV318 cell lines (Sigma) were cultured in DMEM (Corning) supplemented with 10% fetal bovine serum (FBS, Invitrogen), 2 mM glutamine (Life Technologies), and 1% penicillin-streptomycin (pen-step, Life Technologies). KURAMOCHI and OVSAHO (Japanese Collection of Research Bioresources Cell Bank) and SNU-119 (Korean Cell Line Bank) cell lines were cultured in RPMI-1640 (Hyclone) supplemented with 10% FBS and 1% pen-strep. The OVCAR4 cell line (National Cancer Institute Division of Cancer Treatment and Diagnosis Cell Line Repository) was cultured in RPMI-1640 supplemented with 10% FBS and 1% pen-strep. The OVCAR8 cell line (National Cancer Institute Division of Cancer Treatment and Diagnosis Cell Line Repository) was cultured in DMEM (Corning) supplemented with 10% FBS, and 1% penicillin-streptomycin (pen-step, Life Technologies). The OVCAR3 cell line (American Type Culture Collection) was cultured in DMEM with 10% FBS and 1% pen-strep. Caov3 and OVCAR5 cell lines were generous gifts from Professor Anirban Mitra (Indiana University) and were cultured in DMEM with 10% FBS and 1% pen-strep. UWB1.289, UWB1-SyR12 and UWB1-SyR13 cell lines were cultured in RPMI 1640/Mammary Epithelial Growth Media (1:1; Hyclone/PromoCell) supplemented with 3% FBS, MEGM growth factors and 1% pen-step. Primary hOSE cells (ScienCell) were cultured in Ovarian Epithelial Cell Medium (ScienCell, 7311). HGSC precursor cells or immortalized fallopian tube epithelial (FTE; FT190, FT282-E1 and FT282) cells and were cultured in DMEM-Ham’s F12 50/50 (Corning) supplemented with 2% USG (Pall Corporation) or 10% FBS and 1% pen-strep. IOSE-T (IOSE-21, hOSE immortalized with hTERT) cells (Li et al., 2007) were a generous gift from Professor Francis Balkwill (Cancer Research UK) and were cultured in Medium 199/MCDB105 (1:1, Sigma) supplemented with 15% FBS, 1% pen-strep, 10 ng/mL human epidermal growth factor (Life Technologies), 0.5 μg/mL hydrocortisone (Sigma), 5 μg/mL bovine insulin (Cell Applications), and 34 μg protein/mL bovine pituitary extract (Life Technologies). IOSE-SV (IOSE-121, hOSE immortalized with SV40 Large T antigen) were a generous gift from Professor Nelly Auersperg (University of British Columbia) and were cultured in Medium 199/MCDB105 (1:1) supplemented with 10% FBS and 25 µg/ml gentamicin (Life Technologies). HEK293T cells (American Type Culture Collection) was cultured in DMEM with 10% FBS and 1% pen-strep. U2OS-DR-GFP (282C) cells were a kind gift from Professor Jeremy Stark (City of Hope) and grown in McCoy’s 5A (Corning), 10% FBS and 1% pen-strep (Gunn et al., 2011; Gunn and Stark, 2012). OVCAR8-DR-GFP cells were a kind gift from Professor Larry Karnitz and Professor Scott Kaufmann (Mayo Clinic) and grown in the same conditions as the OVCAR8 parental cells as described above (Huntoon et al., 2013). All cell lines were maintained at 37 °C in a humidified incubator with 5% CO_2_. Cell culture medium was changed every 3-5 days depending on cell density. For routine passage, cells were split at a ratio of 1:3-10 when they reached 85% to 90% confluence. Cell lines were authenticated by short tandem repeat (STR) analysis at the DNA Services Facility, University of Illinois at Chicago, and confirmed to be Mycoplasma free by PCR at the Epigenetics Core Facility, University of Nebraska Medical Center. Doxycycline inducible cells were treated every 48 hours with doxycycline (Sigma, solubilized in water) unless otherwise noted.

### Pharmacologic Inhibitors

Cells were treated with the following inhibitors: PARPi (olaparib, ABT-888, solubilized in DMSO) from SelleckChem, ATRi (VE-822, VX970, solubilized in DMSO) from SelleckChem, hydroxyurea (HU, solubilized in water) from Sigma and etoposide (Etop, E1383, solubilized in DMSO) from Sigma.

### Bulk RNA Sequencing (RNA-seq) Analysis

OVCAR8 and Caov3 cells expressing doxycycline inducible shRNAs were seeded on day 0 at ∼50% confluency, respectively, then grown in the presence of 1 μg/ml doxycycline for 48 hours. RNA was harvested on day 2 at ∼80% confluency. DNA-free total RNA was isolated from cells lines using TRIzol (Invitrogen) and Direct-zol RNA Purification Kit (Zymo Research) with in column DNase treatment. The RNA samples were transported to the UNMC DNA Sequencing Core Facility. RNA samples were analyzed with respect to purity and potential degradation. Purity and concentration were assessed by measurement of the A260/280 ratios using a Nanodrop instrument (Thermo Scientific) instrument and only those samples with values of 1.8 to 2.0 underwent further processing. Potential degradation of the samples was assessed by analysis of 200 ng of the RNA with an Advanced Analytical Technical Instruments Fragment Analyzer (Agilent Technologies) and only intact RNA samples were used to generate sequencing libraries. Sequencing libraries were generated by the UNMC NGS Core beginning with 1 µg of total RNA from each sample using the TruSeq V2 RNA sequencing library kit from Illumina following recommended procedures (Illumina Inc., San Diego, CA). Resultant libraries were assessed for size of insert by analysis of an aliquot of each library on a Bioanalyzer instrument (Agilent Technologies). Each library had a unique indexing identifier barcode allowing the individual libraries to be multiplexed together for efficient sequencing. Multiplexed libraries (12 samples per pool) were sequenced across 2 lanes of the HiSeq 2500 DNA Analyzer (Illumina) to generate a total of approximately 20 million 75 bp single reads for each sample. During sequencing the quality was continually monitored regarding cluster number and fluorescence intensity and percentages of reads passing filter with a Q30 score. Following sequencing, samples were demultiplexed to produce FASTQ files. The UNMC Epigenomics Core Facility processed the resulting sequence files based on the following steps. Adaptor sequences and low quality (Phred score: 20) ends were trimmed from sequences using the Trim Galore software package (http://www.bioinformatics.babraham.ac.uk/projects/trim_galore/). Resulting FASTQ files were aligned to the human genome (NCBI37/hg19) using the software TopHat (v2.0.8) (http://ccb.jhu.edu/software/tophat/index.shtml). The software Cufflinks (v2.1.1) http://cole-trapnell-lab.github.io/cufflinks/ was used to estimate the expression values, and Cuffdiff (v2.1.1) was used to determine differential expression. Geneset enrichment analysis (GSEA) was performed using the GSEA software version 3, build 0160, available from the BROAD Institute (http://www.broadinstitute.org/gsea/index.jsp) (Subramanian et al., 2005). Bulk RNA-seq datasets, with genes ranked by log2 fold-change, were compared against the Hallmark Signature genesets (http://software.broadinstitute.org/gsea/msigdb/genesets.jsp?collection=H). Output data included normalized enrichment score (NES) and FDR (q-value). Statistical significance was set at an FDR < 0.25. NES was used for comparing enrichment because it accounts for differences in gene set size and in correlations between gene sets and the expression dataset.

### Single-Cell RNA Sequencing (scRNA-seq) Analysis

FT282 and OVCAR8 cells were seeded on day 0 at ∼50% and ∼30% confluency, respectively, then harvested on day 2 at ∼80% confluence. Cells were trypsinized and suspensions were pipetted several times and passed through a 40 μm cell strainer to ensure a single cell suspension. Cell viability (Trypan blue exclusion) was confirmed to be greater than 90% using the TC20 Automated Cell Counter (Bio-Rad). Cell pellets were washed twice in PBS containing 0.04% BSA, then resuspended in PBS and 0.04% BSA to ∼500,000 cells/ml and transported to the UNMC DNA Sequencing Core Facility for single cell capture, library preparation and scRNA-sequencing. As per manufacturer instructions, approximately ∼2,750 cells were loaded per channel to achieve a target of ∼1,500 captured cells. Single cells were loaded on a Chromium Single Cell Instrument (10x Genomics) to generate single cell Gel Bead in Emulsions (GEMs) for the partitioning of samples and reagents into droplets. GEMs contain oligos, lysed cell components and Master Mix. GEMs were processed as following: briefly, cells were lysed, RNA extracted, and reverse transcribed to generate full-length, barcoded cDNA from the poly A-tailed mRNA transcripts. Barcoded cDNA molecules from every cell were PCR-amplified in bulk followed by enzymatic fragmentation. The Qubit was then used to measure and optimize the insert size of the double-stranded cDNA prior to library construction. Single cell cDNA and RNA-Seq libraries were prepared using the Chromium Single Cell 3’ Library and Gel Bead Kit v2 (10x Genomics) and the products were quantified on the 2100 Bioanalyzer DNA High Sensitivity Chip (Agilent). Each fragment contains the 10x Barcode, UMI and cDNA insert sequence used in data analysis. During library construction Read 2 is added by Adapter ligation. Both single cell libraries were sequenced using the Illumina NextSeq 550 system using the following parameters: pair-end sequencing with single indexing, 26 cycles for Read1 and 98 cycles for Read2. FT282 cells produced a total of 85.6 million reads and OVCAR8 produced 82 million reads, and both had an overall Q30 for the run of 87.6%. Sequencing data were transferred to the Bioinformatics and Systems Biology Core Facility for analysis. The Cell Ranger Single Cell Software (10X Genomics) was used to process raw bcl files to perform sample demultiplexing, barcode processing and single cell 3’ gene counting (https://software.10xgenomics.com/single-cell/overview/welcome). Each dataset was aligned to a combined human (hg19) reference and only cells that were identified as aligned to human in the “filtered” output of the Cell Ranger count module were used in the analysis.

### Statistical Impute of Transcript Dropouts in Single-Cell RNA Sequencing Datasets

The statistical method scImpute was used to accurately and robustly impute the transcript dropouts (zero values) that exist in scRNA-seq data from sequencing small amounts of RNA. The scImpute R package was downloaded from https://github.com/Vivianstats/scImpute and was run in R with default settings. FT282 and OVCAR8 scRNA-seq raw gene expression matrix files (.mtx) were converted into dense expression matrices using ‘cellRanger mat2csv’ command. The scImpute input data consisted of a scRNA-seq matrices with rows representing genes and columns representing cells, and output data consisted of an imputed count matrix with the same dimension.

### Cell Viability Measurements

Cells were seeded at a density of 500-1,000 cells per well in quadruplicate into sterile 96-well plates and treated with the indicated drug for 96 hours. AlamarBlue (BioRad) was used to assess cell viability. Background values from empty wells were subtracted and data were normalized to vehicle-treated control.

### Homologous Recombination (HR) Reporter Assay

U2OS and OVCAR8 cells were previously generated for stable integration of DR-GFP, an HR substrate that generates a functional green fluorescent protein (GFP) upon successful HR after I-SceI cutting (Gunn and Stark, 2012; Huntoon et al., 2013; Pierce et al., 1999). U2OS and OVCAR8 cells were grown in the presence of doxycycline for 48 hours to induce shRNA expression. U2OS cells were transfected with pCBASceI (Addgene; #26477) in the presence of 1 µg/ml doxycycline. Media containing doxycycline was changed 3 hours post transfection and cells were kept in culture for an additional 48 hours. OVCAR8 cells were transduced for 8 hours with AdNGUS24i, an adenovirus expressing I-SceI (Drs. Frank Graham and Phillip Ng). Cells were harvested and fixed with 10% formaldehyde, washed and analyzed by FACS to determine the fraction of GFP positivity.

### *In Vitro* Clonogenic Survival Assay

To assess colony formation, cells were trypsinized, counted and seeded at a density of 500-1,000 cells per well in 6-well dishes in single-cell suspension and allowed to form colonies for 8-12 days. Following incubation, cells were simultaneously fixed and stained in PBS containing 10% methanol and 0.5% crystal violet for 30 minutes at room temperature, rinsed with water and air dried overnight. Colonies containing over 50 cells were manually counted with an inverted light microscope.

### Comet Assay

The Comet Assay Kit (Trevigen) was used according to the manufacturer’s instructions. Briefly, cells were suspended in low melt agarose, layered onto treated slides to promote attachment (500 cells per slide), lysed, and subjected to electrophoresis (1 V/cm) under alkaline conditions to reveal both single- and double-stranded DNA breaks, using the Comet Assay Electrophoresis System II (Trevigen). Samples were then fixed, dried, and stained with SYBR Gold. Images were acquired with a 10X objective lens using the EVOS FL Cell Imaging System (ThermoFisher). Comet tail size was quantified using Comet Analysis Software (Trevigen).

### Flow Cytometry

Cells were fixed in 70% ethanol overnight for cell cycle and γ-H2AX expression analysis. Fixed cells were washed with PBS and incubated overnight in PBS containing 1% BSA, 10% goat serum and pS139-H2AX antibody (Millipore), washed and incubated in goat anti-mouse Alexa Fluor 647 antibody for 30 minutes at RT. Cells were incubated in 50 μg/mL propidium iodide and 100 μg/mL RNase A for 30 minutes, and 10,000 cells per sample were analyzed on a BD FACSarray (BD Biosciences) using 532- and 635-nm excitations and collecting fluorescent emissions with filters at 585/42 nm and 661/16 nm (yellow and red parameters, respectively).

### Production of Lentiviral Particles and *In Vitro* Lentiviral Transduction

Replication-deficient lentivirus was produced by transient transfection of 6.0 μg psPAX2 (Addgene #12260), 2.0 μg pMD2.G (Addgene #12259), and 8.0 μg transfer plasmid into HEK293T cells in a 10 cm dish with Lipofectamine 2000 reagent (Life Technologies), according to the manufacturer’s instructions. Viral supernatants were collected at 48 hours and passed through a 0.2 μm filter. Functional titration was performed by transduction of cells with serially diluted virus in the presence of polybrene (4 μg/ml, Sigma) for 6 hours followed by puromycin (Life Technologies) selection 48 hours post-infection.

### RNA Interference and Plasmid Constructs

**Table S1** contains a list of all siRNAs and shRNAs. **Table S2** contains a list of all plasmids. All cloning was sequence verified. Plasmids were transfected with Lipofectamine 2000 (Life Technologies), according to the manufacturer’s instructions. siRNAs were transfected with Lipofectamine RNAiMax reagent (Life Technologies), according to the manufacturer’s instructions.

### CRISPR Knockout

CRISPR-mediated knockout was performed as previously described (Ran et al., 2013). Briefly, Lipofectamine 2000 (Life Technologies) was used to transfect HEK293T cells with PX458 (Addgene #48138), which expresses Cas9 and a guide RNA. **Table S1** contains a list of all sgRNAs. Guide RNAs were designed using the Genetic Perturbation Platform Web Portal (https://portals.broadinstitute.org/gpp/public/) and the highest scoring guide RNAs were then empirically tested for their cutting efficiency. Isolation of clonal cell lines was achieved with fluorescence-activated cell sorting (FACS) following by plating of GFP positive cells into 96-well dishes for clonal growth. DNA was isolated from clones with QuickExtract (Epicentre) for genotyping and replica plated for clonal expansion. Genotyping was performed by PCR with annealing temperature of 60 °C and 30 cycles for all primer pairs listed in Table S1. Primers were designed with NCBI Primer-BLAST then empirically tested by gradient end-point PCR to optimize the specificity and sensitivity.

### Reverse Transcription Quantitative PCR (RT-qPCR)

Total RNA was purified using TRIzol (Invitrogen) and RNA was DNase-treated using the DNA-free kit (Ambion) or Direct-zol RNA Purification Kit (Zymo Research) with in column DNase treatment. RNA quality was determined by RNA denaturing gel. DNase-treated RNA was converted to cDNA using the iScript cDNA Synthesis Kit (BioRad). One µl of 1:5 cDNA sample dilutions were used for qPCR reactions. Standard curves were prepared using gel-purified end-point RT-PCR products. All samples were run in triplicate using the CFX Connect Real-Time System (Bio-Rad) and all gene expression data were normalized to 18s rRNA. PCR was performed with an annealing temperature of 60 °C and a total of 45 cycles for all primer pairs. Dissociation curves were performed to confirm specific product amplification. RT-qPCR standards for each gene were generated from a mixture of human or mouse cell cDNA via end-point RT-PCR then gel purification, using the appropriate primer pair. Gradient PCR reactions were performed with the C1000 Touch Thermal cycler (Bio-Rad) to determine the annealing temperatures for each primer set. Primer sequences are listed in **Table S1**. Primer sequences corresponding to each gene for the mRNA expression analysis were designed using NCBI Primer Blast (Ye et al., 2012) or selected from those previously reported in the literature.

### Western Blot

Whole cell protein extracts were prepared with RIPA buffer [1X PBS, 1% NP40, 0.5% sodium deoxycholate, 0.1% sodium dodecyl sulfate (SDS)] supplemented with protease and phosphatase inhibitors (Sigma) and centrifuged at 4 °C for 10 minutes at 14,000 *g*. Nuclear extracts were prepared using the NE-PER Nuclear and Cytoplasmic Extraction Kit (Thermo Scientific) supplemented with protease and phosphatase inhibitors. Protein concentration was determined by the BCA protein assay (Thermo Scientific). Equal amounts of protein (30-50 µg) were fractionated on 4-12% gradient SDS-polyacrylamide gel electrophoresis gel (Invitrogen) and transferred to PVDF membrane (Roche). Membranes were stained with Ponceau S to confirm efficient transfer and equal loading then blocked with 5% nonfat dry milk in Tris-buffered saline Tween-20 (TBST) for 1 hour at room temperature. The membranes were incubated with primary antibodies in 5% nonfat dry milk or 5% BSA in TBST at 4°C overnight followed by incubation with secondary antibody in 5% nonfat dry milk in TBST for 1 hour at room temperature. Primary antibodies are listed in **Table S3**. Enhanced chemiluminescence (Thermo Scientific) was used for protein detection. Quantification of protein expression was performed using ImageJ software (Image Processing and Analysis in Java, National Institute of Health) (Schneider et al., 2012).

### Co-Immunoprecipitation

Cells expressing empty vector, and Flag- or HA-tagged ORFs were lysed with M-PER (Pierce) containing Halt Protease and Phosphatase Cocktail (Pierce), and Nuclease S1. Lysates were mixed end over end at 4 °C for 30 minutes then centrifuged at 4 °C for 10 minutes at 14,000 *g* to eliminate cellular debris. Protein concentration was determined by the BCA protein assay (Thermo Scientific). Immunoprecipitation was performed with anti-Flag or HA magnetic beads and 500 µg total protein. Samples were incubated overnight at 4 °C with end over end mixing. The next day the magnetics beads were washed, and protein eluted with sample loading buffer. The entire sample was loaded on the Western blot along with 5% total protein as input.

### Illumina Infinium Human Methylation 450K Datasets

The TCGA pan-cancer DNA methylation (HumanMethylation450K) dataset was downloaded from UCSC Xena (http://xena.ucsc.edu), consisting of 9,753 samples. The TCGA pan-cancer dataset contains the DNA methylation 450K array beta values, compiled by combining available data from all TCGA cohorts. The GDSC1000 collection (The Genomics of Drug Sensitivity in Cancer Project, Welcome Sanger Institute) was downloaded from NCBI GEO (GSE68379) (Iorio et al., 2016). The NCI-60 cell line DNA methylation 450K dataset (NCBI GEO: GSE66872) was downloaded from CellMiner (Reinhold et al., 2012; Shankavaram et al., 2009). For all datasets, the DNA methylation profile was measured experimentally using the Illumina Infinium HumanMethylation450 platform and Infinium HumanMethylation450 BeadChip Kit. The method consists of approximately 450,000 probes querying the methylation status of CpG sites within and outside CpG islands. The DNA methylation score of each CpG is represented as a beta value, which is a normalized value between 0 (fully-unmethylated) and 1 (fully-methylated).

### Sodium Bisulfite DNA Sequencing

Sodium bisulfite DNA sequencing was used to determine the methylation status of the F/R-BDP region (Clark et al., 1994). Sanger sequencing analysis was performed by the University of Nebraska Medical Center DNA Sequencing Core Facility. Genomic DNA was isolated using the Puregene Tissue Kit (Qiagen), and then chemically converted with sodium bisulfite using the EZ DNA Methylation Kit (Zymo Research Corp.). The F/R-BDP region containing a CpG island was amplified from the bisulfite-converted DNA using methylation PCR specific primers, designed using MethPrimer (**Table S1**) (Li and Dahiya, 2002). Gradient PCR reactions were analyzed using a C1000 Touch Thermal cycler (Bio-Rad) to optimize annealing temperatures for the primer set. After gel purification using the QIAquick Gel Extraction Kit (QIAGEN), PCR products were cloned using either the TOPO TA Cloning Kit (Invitrogen). Between 9 and 15 individual clones were sequenced for each sample. DNA sequence information was analyzed using the Lasergene SeqMan Pro program (DNASTAR, Inc.).

### 5’ RNA Ligase-Mediated Rapid Amplification of cDNA Ends (RLM-RACE) Mapping

The transcriptional start site of *FOXM1* and *RHNO1* in different cell lines was determined using the FirstChoice RLM-RACE Kit (Ambion), according to the manufacturer’s instructions. RNA for RLM-RACE analysis was isolated using TRIzol reagent (Invitrogen) followed by purification with Direct-zol RNA MiniPrep with in column genomic DNA digestion. RLM-RACE assay allows for the specific amplification of 5’ capped RNA, which is found only on full-length mRNA transcripts. The specific outer and inner (nested) primers in combination with adaptor-specific primers were used for amplifying the 5’ ends of *FOXM1* and *RHNO1* mRNA. The 5’ end specific primers are listed in **Table S1**. As negative controls, non-tobacco alkaline phosphatase (TAP)- treated aliquots from each RNA source were utilized; in all cases, this did not yield any specific product amplification (data not shown). In contrast, FOXM1- and RHNO1-specific PCR products of various sizes were yielded from TAP-containing reactions. PCR products were separated on 2% agarose gels, excised, and subsequently purified using the QIAquick gel extraction kit (Qiagen). Gel-purified PCR products were cloned using the TOPO TA Cloning Kit (Invitrogen), and individual clones were sequenced using standard methods.

### Promoter Activity Luciferase Reporter Assay

The 6X-FOXM1 luciferase reporter assay was performed with pGL4-6X-FOXM1 (firefly) and pRL-TK (renilla, transfection control). The FOXM1 promoter reporter assay was performed with PGL3-FOXM1 promoter (−1.1 kb) and PGL3-FOXM1 promoter (−2.4 kb) and SEAP-PGK (transfection control). Firefly and Renilla luciferase activities were measured 24 hours after transfection using the Dual-Luciferase Reporter Assay System (Promega) and GloMax 20/20 luminometer (Promega). Firefly and renilla luciferase activities were expressed in arbitrary units as displayed on the luminometer after a 10 second integration time. SEAP activity was measured 24 hours after transfection using the Phospha-Light kit (Thermo Scientific) and POLARstar OPTIMA microplate reader (BMG Labtech). All transfections were performed in triplicate within each individual experiment.

### University of California Santa Cruz (UCSC) Genome Browser Data Retrieval

The genomic region of *FOXM1* and *RHNO1* was retrieved from UCSC Genome Browser (http://genome.ucsc.edu) (Kent et al., 2002) using the human genome build hg19 with genomic coordinates chr12:2,966,265-2,999,264. The following tracks were selected and displayed in the Genome Browser: *FOXM1* and *RHNO1* mRNA, CpG island, Encode E2F1, H3K4Me3 and H3K27Ac ChIP-seq tracks, and conserved genome tracks from 100 vertebrates and mammalian genomes. The Genome Browser screen shot tool was used to obtain images.

### The Cancer Genome Atlas (TCGA), Cancer Cell Line Encyclopedia (CCLE), Genotype-Tissue Expression (GTEx), and RIKEN Fantom5 Datasets

TCGA, CCLE and GTEx genomic profiled datasets were retrieved from cBioPortal or UCSC Xena Browser. The genomic profile of *FOXM1* and *RHNO1* were analyzed in the HGSC (Ovarian Serous Cystadenocarcinoma-TCGA Provisional) dataset for putative somatic copy-number alterations from GISTIC (Beroukhim et al.) using Onco Query Language (OQL) and mRNA expression (RNA seq V2 RSEM). TOIL GTEx (cell lines removed) and TCGA HGSC RNA-seq datasets were obtained from UCSC Xena. CAGE (Cap Analysis of Gene Expression) analysis of mouse tissues in RIKEN FANTOM5 project (RNA-seq) dataset was retrieved from www.ebi.ac.uk.

### *FOXM1* and *RHNO1* Correlations in Publically-Available Single-Cell RNA Sequencing Datasets

Single-cell RNA-sequencing datasets from mouse normal colon epithelium (GSE92332) (Haber et al., 2017) and human melanoma (GSE72056) (Tirosh et al., 2016) were downloaded from NCBI GEO. Single-cell RNA sequencing data for human high-grade serous ovarian cancer tissue was obtained from a published dataset (Winterhoff et al., 2017). Single-cell RNA sequencing data were normalized with sclmpute (Li and Li, 2018).

### Statistical Analyses

Student’s t-test was used to compare differences between means of two groups. Mann-Whitney test was used to compare differences between medians of two groups. One-way analysis of variance (ANOVA) with a post-test for linear trend was used to compare two or more groups. For all analyses, significance was inferred at P < 0.05 and P values were two-sided. GraphPad Prism statistical software (GraphPad Software, Inc) was used for statistical analysis.

### Data Deposit

Upon publication, bulk and scRNA sequencing data will be deposited into GEO.

## Acknowledgments

We thank Dr. Keith Johnson for key initial insight, and Dr. Gargi Ghosal for critical reading of the manuscript. We thank Dr. Anirban Mitra for Caov3 cells, Drs. Larry Karnitz and Scott Kaufmann for OVCAR8 DR-GFP cells, Drs. Jeremy Stark and Tadayoshi Bessho for U2OS DR-GFP cells, Drs. Frank Graham and Philip Ng for adenovirus expressing SceI, Dr. Stephen Elledge for mutant RHNO1 constructs, and Dr. Aziz Sancar for RHNO1 antibody. We thank the UNMC Epigenomics, Genomics, and Flow Cytometry Core Facilities for support. This work was funded by a Rivkin Center for Ovarian Cancer Bridge Award, The Fred & Pamela Buffett Cancer Center, NCI P30 CA036727, NIH T32CA009476, NIH/NCI F99CA212470, and a UNMC Program of Excellence Assistantship.

## Author Contributions

Conceptualization, CJB, A.R.K; Methodology, CJB; Formal Analysis, M.A., D.K.; Investigation, CJB, C.B., L.C.; Resources, R.D., K.O, L.Z., Data Curation, M.A., D.K.; Writing-Original Draft; CJB and A.R.K.; Writing-Review & Editing, CJB, A.R.K.; Supervision, A.R.K.; Funding Acquisition, CJB, A.R.K.

---

**Table S1. Oligonucleotides.**

**Table S2. Plasmids.**

**Table S3. Antibodies.**

## Supplementary Figure Legends

**Figure S1. Recurrent sites of DNA copy number amplification in HGSC determined by GISTIC.** The GISTIC plot shows 33 recurrent sites of DNA copy number amplification in 579 TCGA HGSC samples. The x-axis displays statistical significance as FDR q-value. The black vertical line indicates the threshold for significance, which is set with an FDR q-value less than 0.25. Chromosomes listed on the left are arranged from chromosome 1 on top to chromosome X on the bottom. Significantly amplified cytobands are listed on the right. 12p13.33 peak is highlighted on the plot. The 33 genes contained in the 12p13.33 amplicon are shown to the right. The GISTIC data was retrieved on 8/21/2015 from the Broad Institute TCGA Genome Data Analysis Center. TCGA HGSC SNP6 Copy number analysis (GISTIC2).

**Figure S2. GTEx 12p13.33 gene expression correlations.** Hierarchical clustering dendogram illustrating the correlation of 33 genes in the TCGA HGSC 12p13.33 amplicon in normal human tissues (GTEx).

**Figure S3.** UCSC Genome Browser (http://genome.ucsc.edu, human genome build hg19 with genomic coordinates chr12:2,966,265-2,999,264) displaying *FOXM1* and *RHNO1* genes. The following tracks are displayed in the browser: *FOXM1* and *RHNO1* mRNA, CpG island, Encode E2F1, H3K4Me3 and H3K27Ac ChIP-seq track, and conserved genome tracks 100 vertebrates and mammalian. The FOXM1RHNO1 promoter is highlighted with the light blue vertical line.

**Figure S4. *FOXM1* and *RHNO1* mRNA expression (RNA-seq) in normal and cancer. A.** *FOXM1* and *RHNO1* expression (RNA-seq) in GTEx and TCGA normal tissues and TCGA tumor tissues. **B**. *FOXM1* and *RHNO1* expression ratio in normal and cancer, and *MKI67* normalized ratio on the far right for TCGA normal and tumor tissues.

**Figure S5. Distribution of DNA methylation at the F/R-BDP in TCGA normal and tumor tissues, and CCLE cancer cell lines.** Beta values represent the level of DNA methylation, averaged across seven CpG sites within the F/R-BDP CGI. A value of one is the highest level of methylation and zero is the lowest level of methylation.

**Figure S6. F/R-BDP region DNA methylation analysis using bisulfite clonal sequencing.** Sodium bisulfite clonal sequencing of the FOXM1/RHNO1 bidirectional promoter region was performed on the indicated samples. The NCBI suggested TSS is indicated by the red broken arrows. The intergenic space spans between the two arrows. Filled and open circles indicate methylated and unmethylated CpG sites, respectively, and each row represents one sequenced allele.

**Figure S7. *FOXM1* and *RHNO1* expression correlations in primary vs. recurrent HGSC.** Two different patient-matched sets of primary vs. recurrent chemoresistant HGSC were used (Kreuzinger et al., 2017; Patch et al., 2015). **A-B)** Expression correlation in (**A**) primary and (**B**) recurrent patients from Kreuzinger et al. **C-D)** Expression correlation in (**C**) primary and (**D**) recurrent patients from Patch et al.

**Figure S8. *FOXM1/RHNO1* expression ratios in primary vs. recurrent HGSC.** Two different patient-matched sets of primary vs. recurrent chemoresistant HGSC were used (Kreuzinger et al., 2017; Patch et al., 2015). Mann-Whitney expression p-value is indicated.

**Figure S9. *FOXM1* and *RHNO1* expression in individual patient-matched primary and recurrent chemoresistant HGSC samples.** The data presented are from Patch et al. (Patch et al., 2015). Patients showing coordinated *FOXM1* and *RHNO1* induction at recurrence are demarcated by asterisks.

**Figure S10. Mapping of the *FOXM1* and *RHNO1* transcriptional start sites (TSS).** RLM-RACE mapping the 5’ end of *FOXM1* and *RHNO1* mRNAs. The red left and right arrows indicate the orientation of each gene. The left y-axis indicates the sequenced clones from the respective HGSC and FTE cell line. NCBI predicted TSS is shown at the bottom. The green line at the top indicates the bidirectional promoter cloned in reporter assays in subsequent experiments. The bottom lines represents a scale in bp length.

**Figure S11. F/R-BDP promoter activity measurements in individual cells. A-D.** Individual cell F/R-BDP reporter activity assay. **A.** F/R-BDP reporter construct with the promoter colored in black. FOXM1 drives GFP and RHNO1 drives RFP. **B-C.** FACS plots of *FOXM1* and *RHNO1* promoter activity in HEK293T cells transfected with empty reporter construct. **D-E.** FACS plots of *FOXM1* and *RHNO1* promoter activity in HEK293T cells transfected with F/R-BDP promoter reporter construct.

**Figure S12. Clonogenic survival for HGSC cells following FOXM1 or RHNO1 CRISPR gene knockout.** OVCAR8 and CAOV3 cells engineered for FOXM1 or RHNO1 CRISPR knockout were seeded for protein or clonogenic survival. **A-B.** Cells were for 72 hours to harvest protein followed by Western blot analysis to confirm knockout efficiency. **C.** Cells were seeded into a 6-well dish, in triplicate, at a density of 500 or 1000 cells, respectively. Media was replenished every 48 hours. Clonogenic survival was measured at 12 and 14 days, respectively, after the cells were fixed with methanol and stained with crystal violet. **D.** Colonies containing more than 50 cells were counted and clonogenic survival was quantified as an average of the replicates. **E.** Total apoptotic cells for OVCAR8 FOXM1 or RHNO1 knockout cells was determined by Annexin V staining and FACS. Etoposide was usd a positive control CTL = non-targeting guide RNA. test *P* value is shown. *P* value designation: **** < 0.0001, *** < 0.001, ** < 0.01, * < 0.05.

**Figure S13. RNA-sequencing of HGSC cells with FOXM1 shRNA knockdown**. CAOV3 and OVCAR8 cells with dox-inducible non-targeting or FOXM1 shRNA were grown in the presence dox for 48 hours and cells were harvested for RNA or protein. Sample were prepared in triplicate for each group. **A.** Western blot analysis of FOXM1 protein expression to confirm knockdown efficiency. **B.** RNA sequencing performance characteristics. **C-G**. GSEA plots, showing normalized enrichment scores (NES), and false discovery rate q values (FDR). **C-D.** GSEA analysis of the top hallmark pathways for OVCAR8 FOXM1 knockdown cells. **E.** HR repair pathway data for OVCAR8 cells. **F-H**. GSEA analysis of the top two hallmark pathways for CAOV3 FOXM1 knockdown cells. **I-K.** RT-qPCR validations of selected FOXM1 target genes in CAOV3 and OVCAR8 cells.

**Figure S14. RHNO1 interactions with 9-1-1 checkpoints proteins in 293T and OVCAR8 cells. A.** Schematic of RHNO1 wild-type and SWV mutant proteins. The proteins are the same molecular weight but they are different for residues 55-61. The residues at 55-61 were all converted to alanine for the SWV mutant thus disrupting its ability to interact with 9-1-1 checkpoint proteins**. B.** 293T were transfected with empty vector, vector expressing HA tagged RHNO1 wild-type or SWV mutant. Protein was harvested 24 hours post transfection for co-immunoprecipitation and Western blot analysis. **C**. OVCAR8 cells were seeded for growth for 24 hours then treated with vehicle or 5 mM HU for 1 hour then cells were harvest for subcellular fraction into soluble nuclear and chromatin bound protein followed by Western blot analysis to determine RHNO1 localization. Histone H3 was used as the chromatin control and Sp1 as the soluble nuclear control. **D.** OVCAR8 inducible RHNO1 knockdown cells were engineered for inducible expression of HA tagged RHNO1 wild-type or SWV mutant and grown in the presence of dox for 72 hours then protein was harvested for co-immunoprecipitation and Western blot analysis. **E.** OVCAR8 Western blot measurement of P-CHEK1 following RHNO1 knockdown. **F.** Quantification of gH2AX positive cells using combined cell cycle analysis and antigen staining by flow cytometry. **G.** COMET analysis of DNA strand breakage in the indicated treatments of OVCAR8 cells.

**Figure S15. RHNO1 interaction with the 9-1-1 checkpoint clamp promotes HGSC cell survival.** OVCAR8 dox-inducible RHNO1 knockdown cells engineered for inducible expression of HA tagged RHNO1 wild-type or SWV mutant were seeded for RNA and protein extractions or clonogenic survival assays. **A-B.** Cells were seeded in the presence of dox and grown for 72 hours to harvest RNA and protein extracts, followed by (**A**) RT-qPCR or (**B**) Western blot to confirm knockdown efficiency**. C.** Cells were seeded into a 6-well dish, in triplicate, in the presence of dox at a density of 500 cells. Media containing dox was replenished every 48 hours. Clonogenic survival was measured at 12 days after the cells were fixed with methanol and stained with crystal violet. **D.** Colonies containing more than 50 cells were counted and clonogenic survival was quantified as an average of the replicates. NS = non-targeting shRNA. Homozygous deletion of RHNO1 in FTE cells was achieved with CRISPR-Cas9 and guide RNAs that targeted near the start and stop codon. **E-F.** RHNO1 homozygous knockout in FT282 cells was confirmed by (**E)** PCR with genomic DNA and (**F**) *RHNO1* mRNA expression by RT-PCR. **G.** FTE parental and RHNO1 clonal knockout cells were seeded in quadruplicate and grown for a period of one-week. Cell viability was measured with the AlamarBlue assay every 24 hours. T-test P value is shown. P value designation: **** < 0.0001, *** < 0.001, ** < 0.01, * < 0.05.

