## Supplemental Figures for "Co-regulation and functional cooperativity of FOXM1 and RHNO1 bidirectional genes in ovarian cancer"

### Figure S1. Barger *et al*.

A

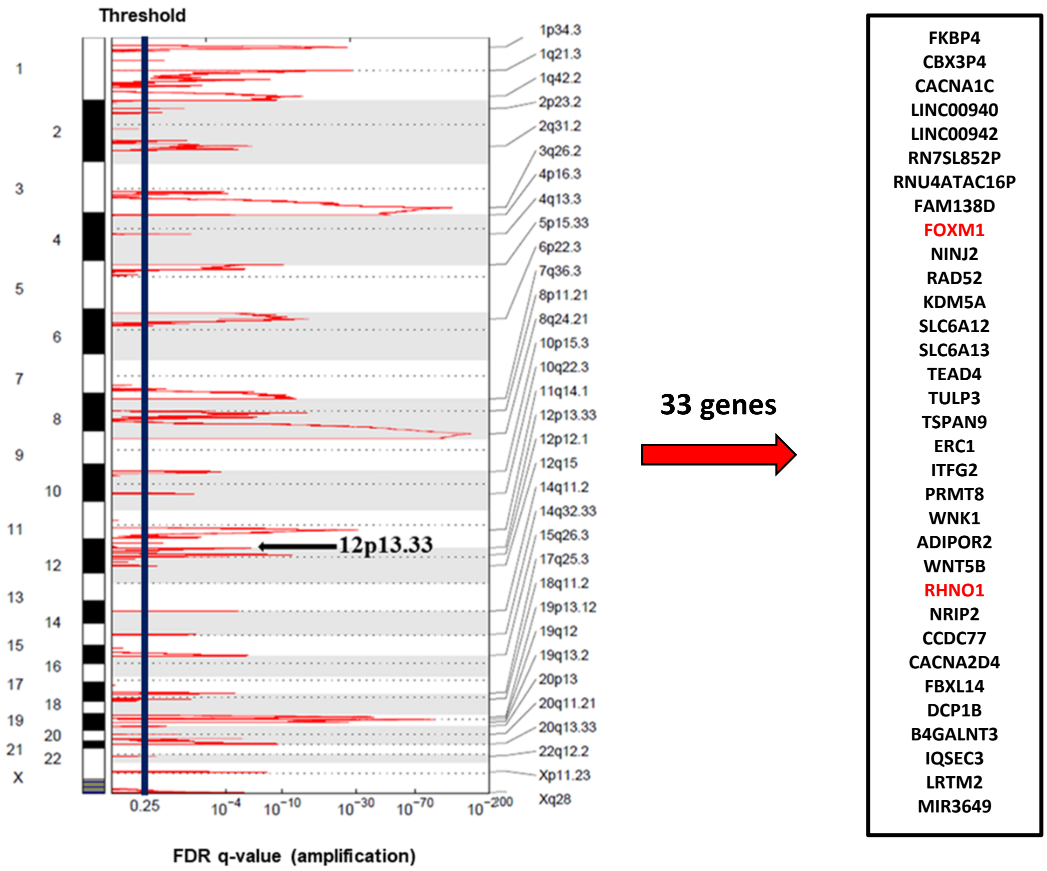

**Figure S2. Barger *et al*.**

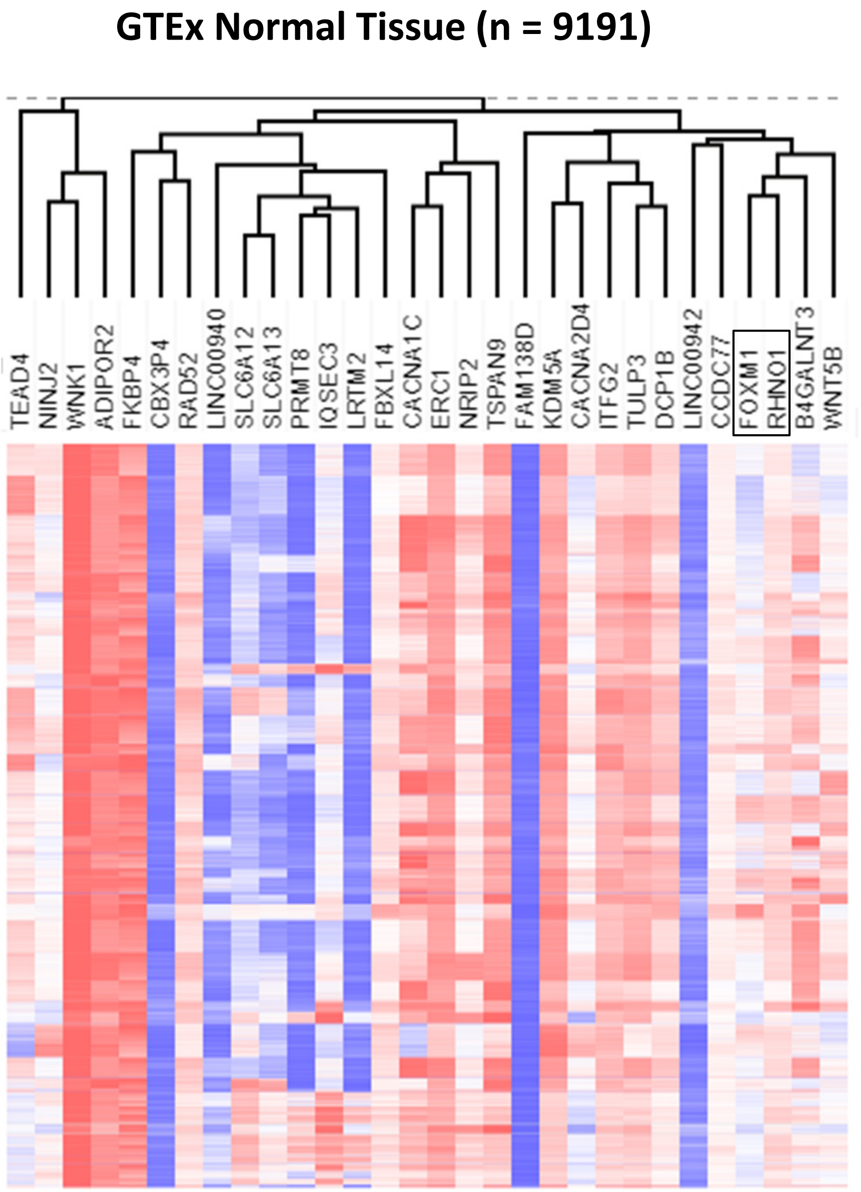

Figure S3. Barger *et al*.

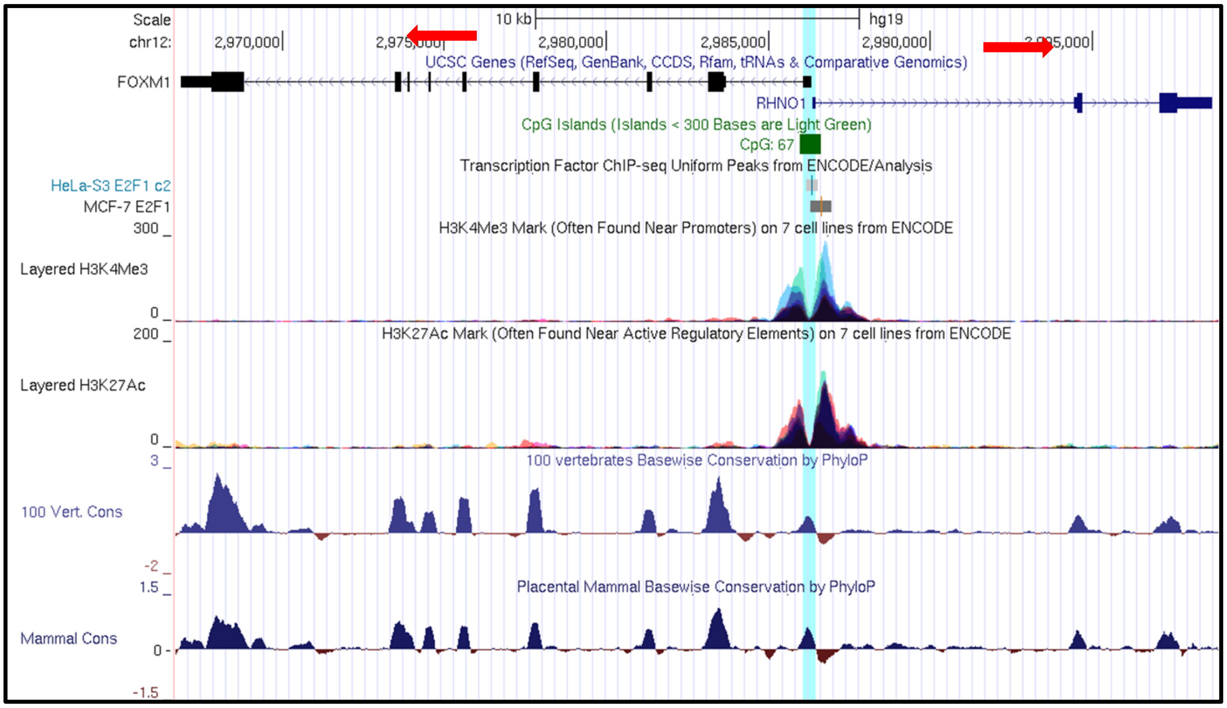

**Figure S4. Barger *et al*.**

A

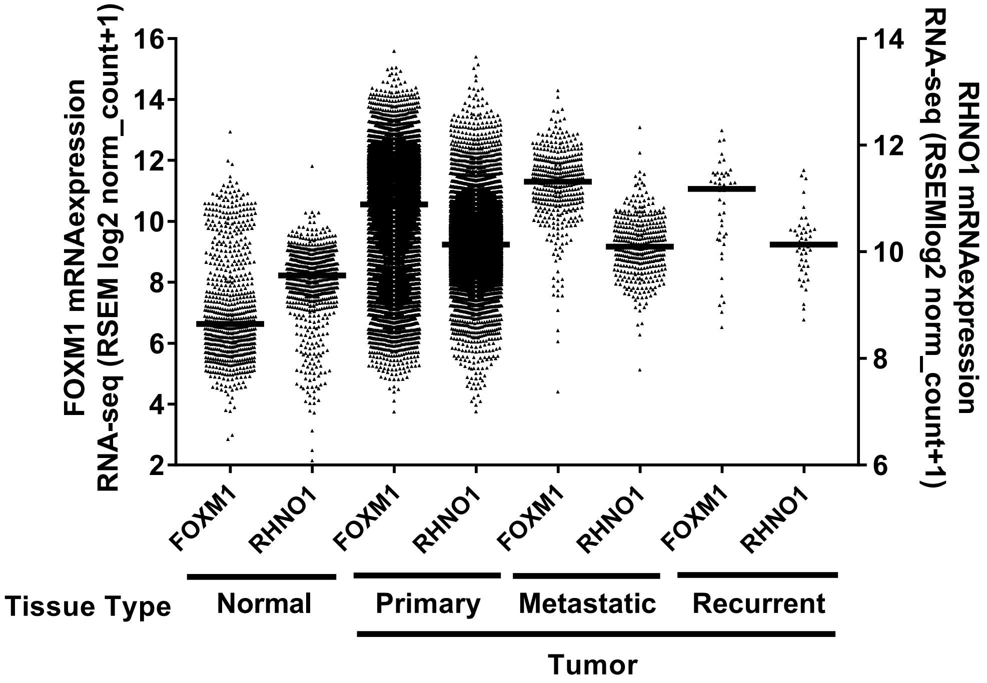


B

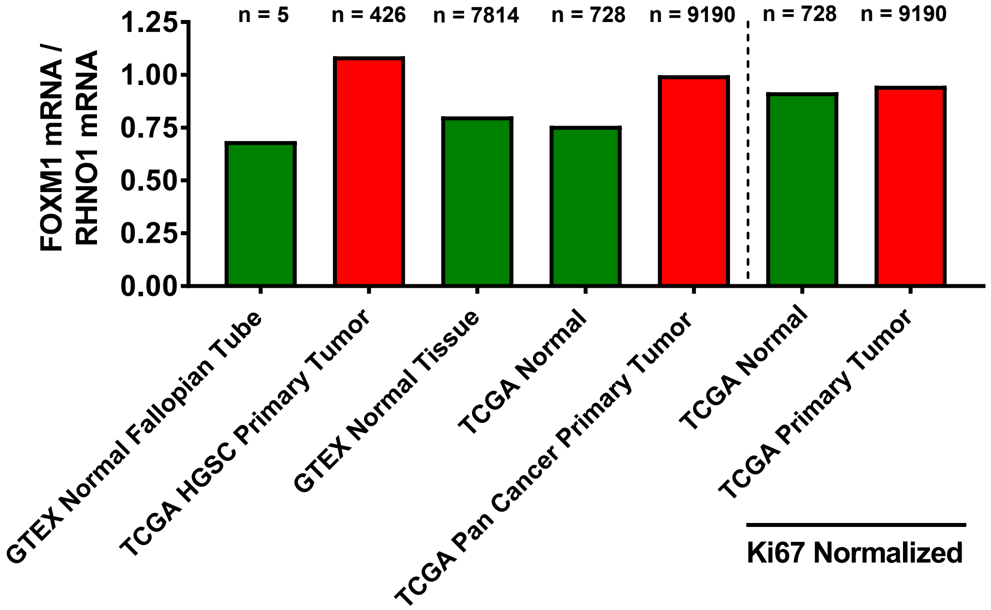

**Figure S5. Barger *et al*.**

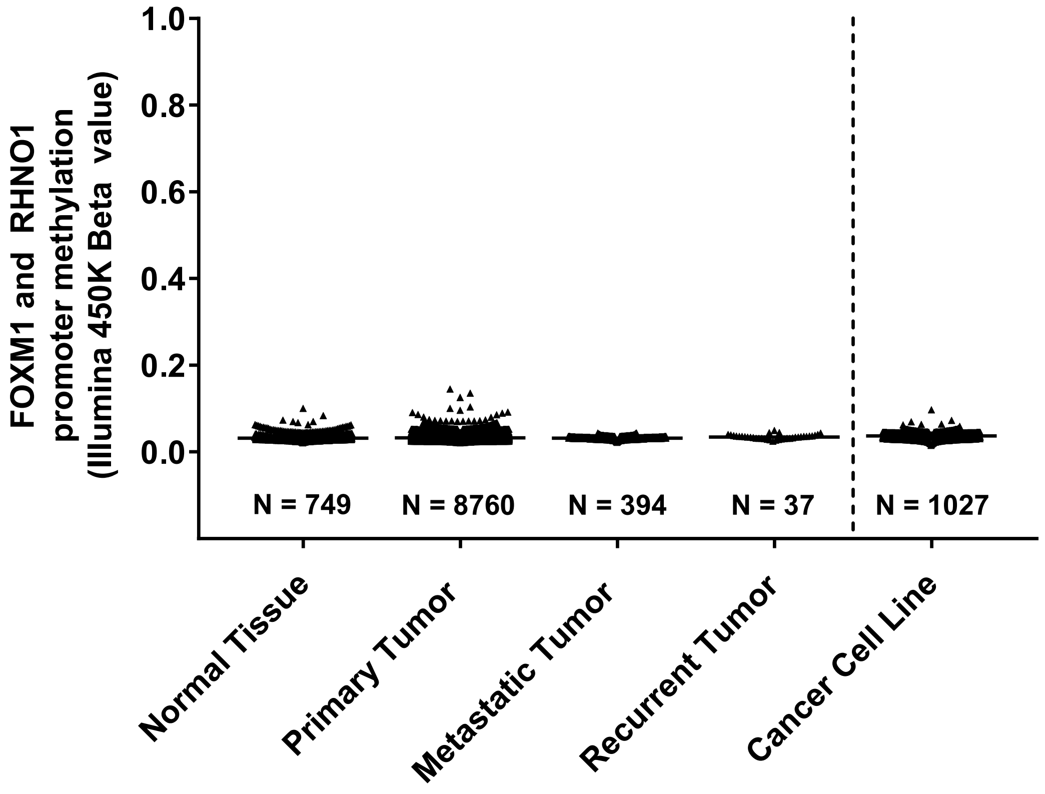

**Figure S6. Barger *et al*.**

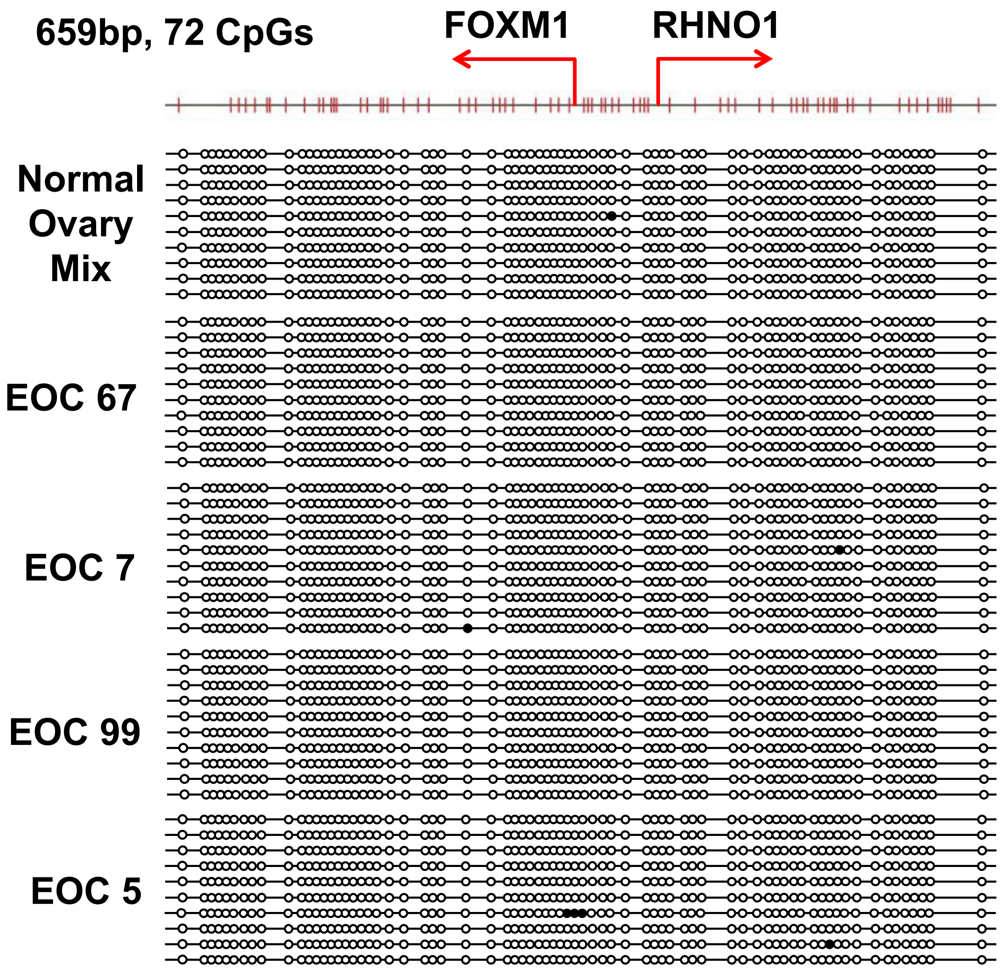

### Figure S7. Barger *et al*.

**Cacsire's Data (left: Primary n=66, right: Recurrent n=66):**

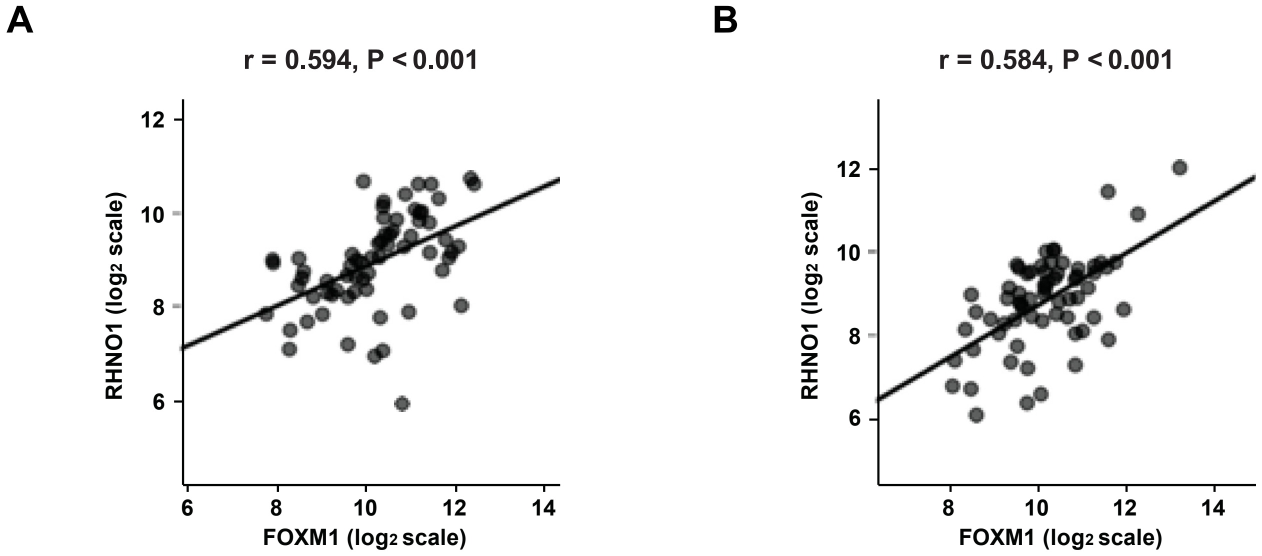


**Patch's Data (left: Primary, n=81, right: Recurrent n=26):**

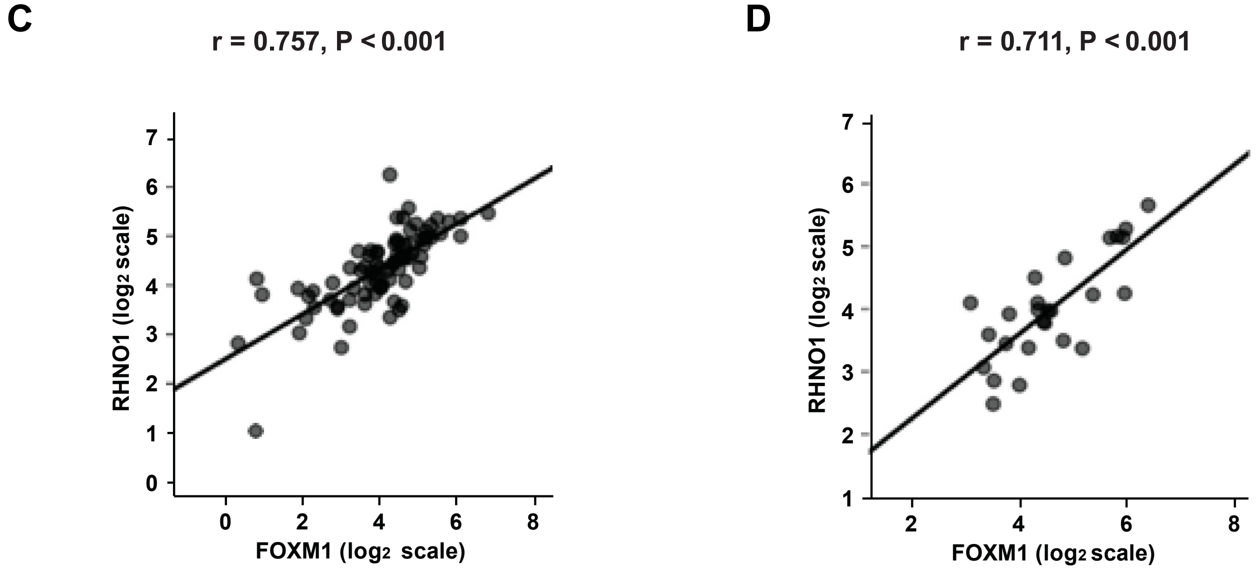

#### Figure S8. Barger *et al*.

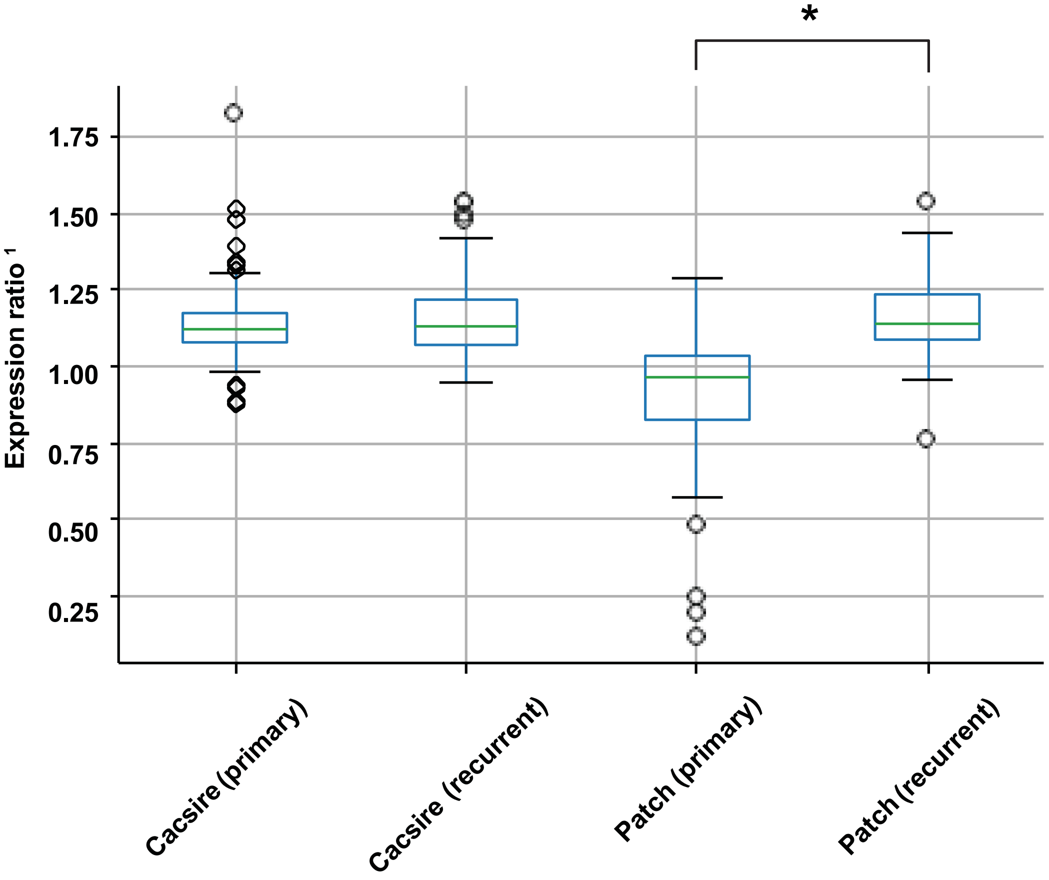


---

[1] $\text{Expression Ratio} = \dfrac{\text{Expression of FOXM1}}{\text{Expression of RHNO1}}$

* Mann-Whitney (U test): P < 0.001

**Figure S9. Barger *et al*.**

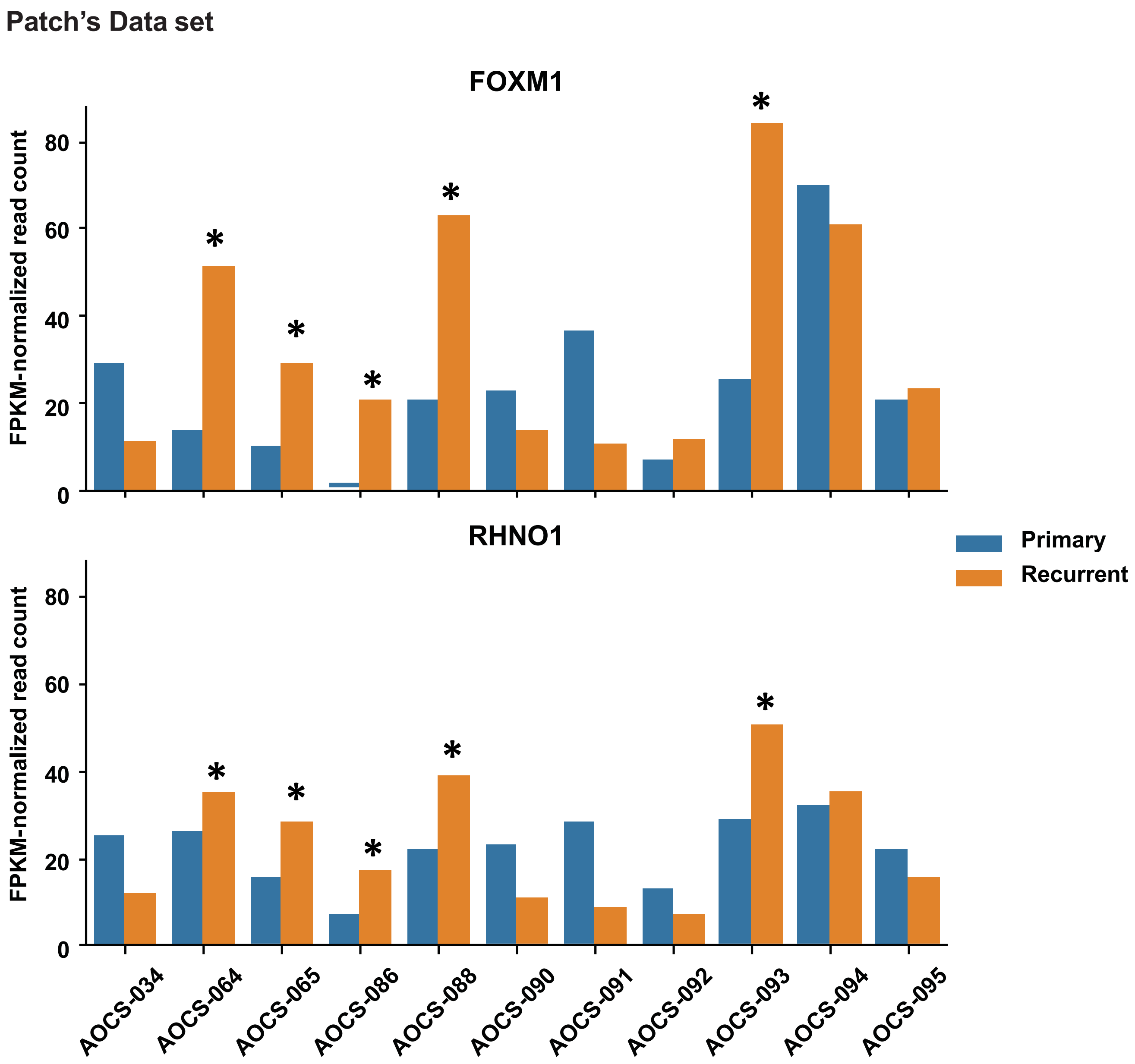

**Figure S10. Barger *et al*.**

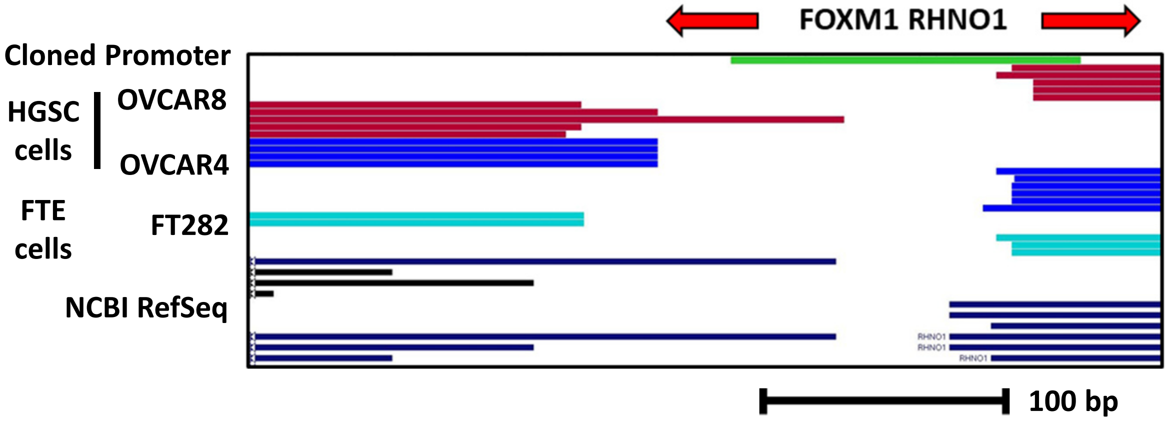

**Figure S11. Barger *et al*.**

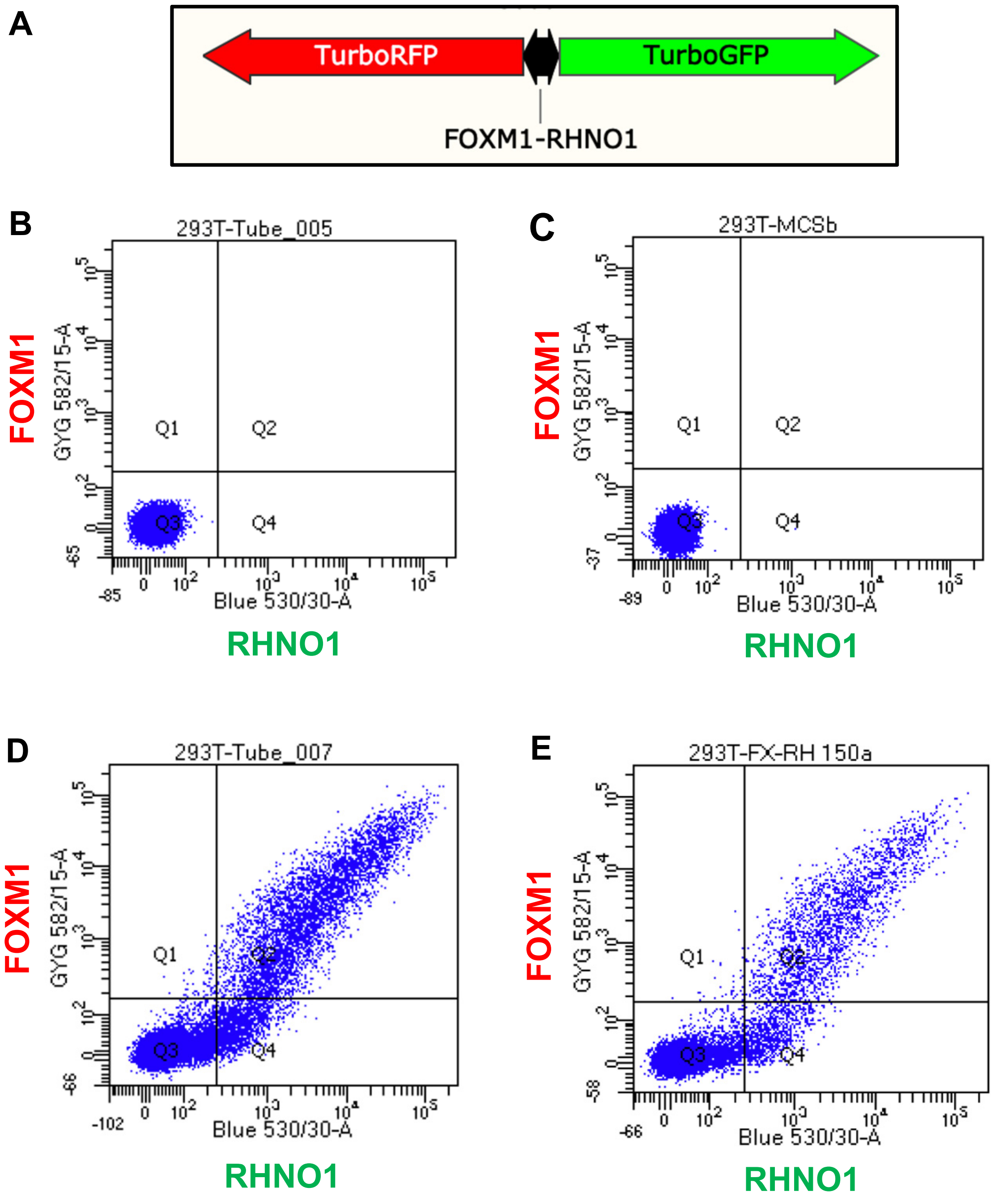

**Figure S12. Barger *et al*.**

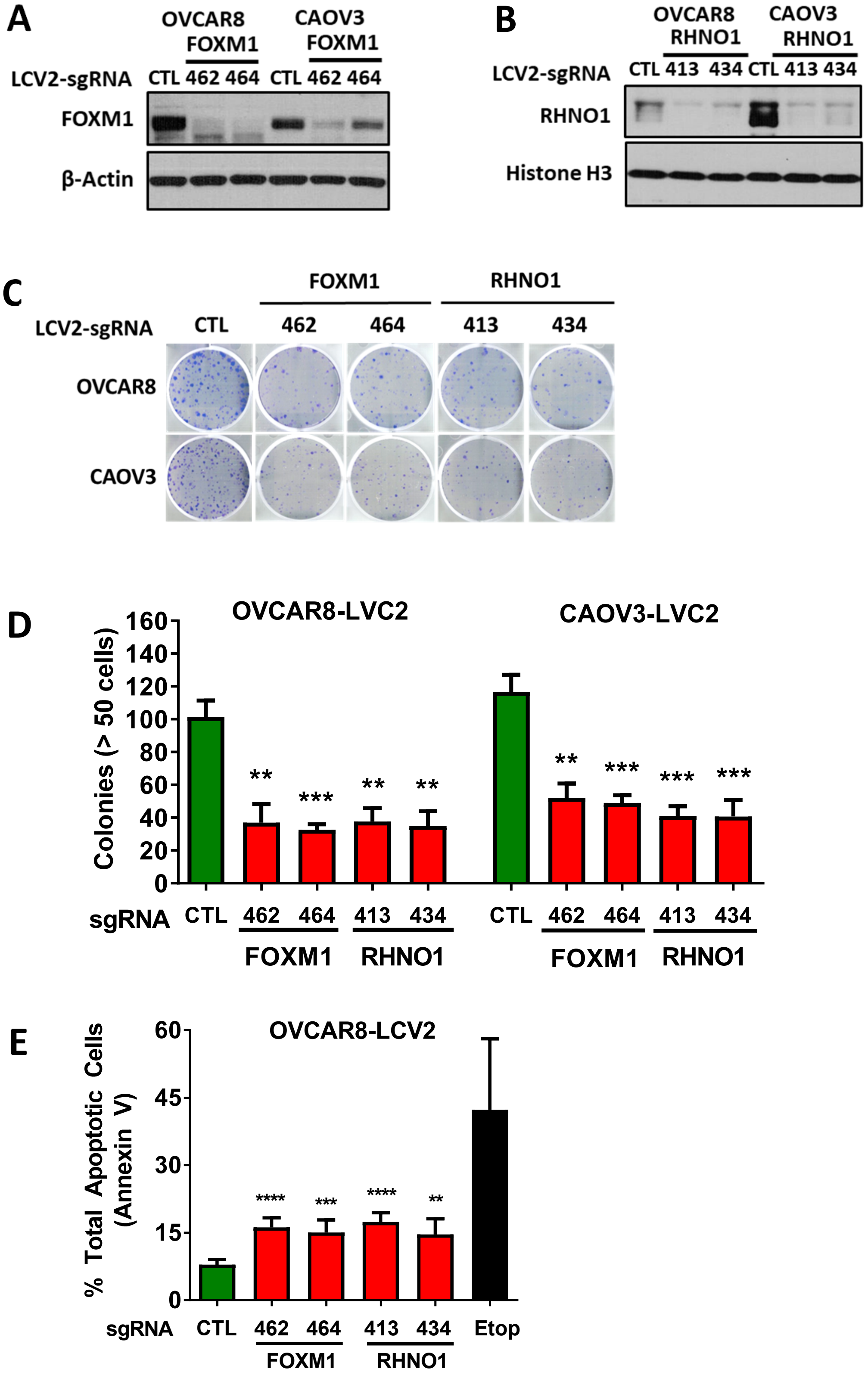

### Figure S13. Barger *et al.*

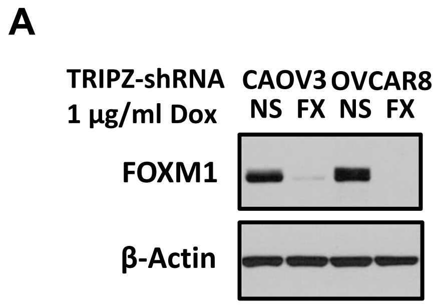


**B**

HiSeq 2500 DNA Analyzer (Illumina)
Triplicate replicates with 75 bp single reads

| Sample name | Means reads ± SD ($10^6$) |
|---|---|
| CAOV3-shNS | 19.7 ± 3.3 |
| CAOV3 -shFOXM1 | 22.1 ± 2.5 |
| OVCAR8-shNS | 21.5 ± 1.9 |
| OVCAR8-shFOXM1 | 21.2 ± 0.8 |

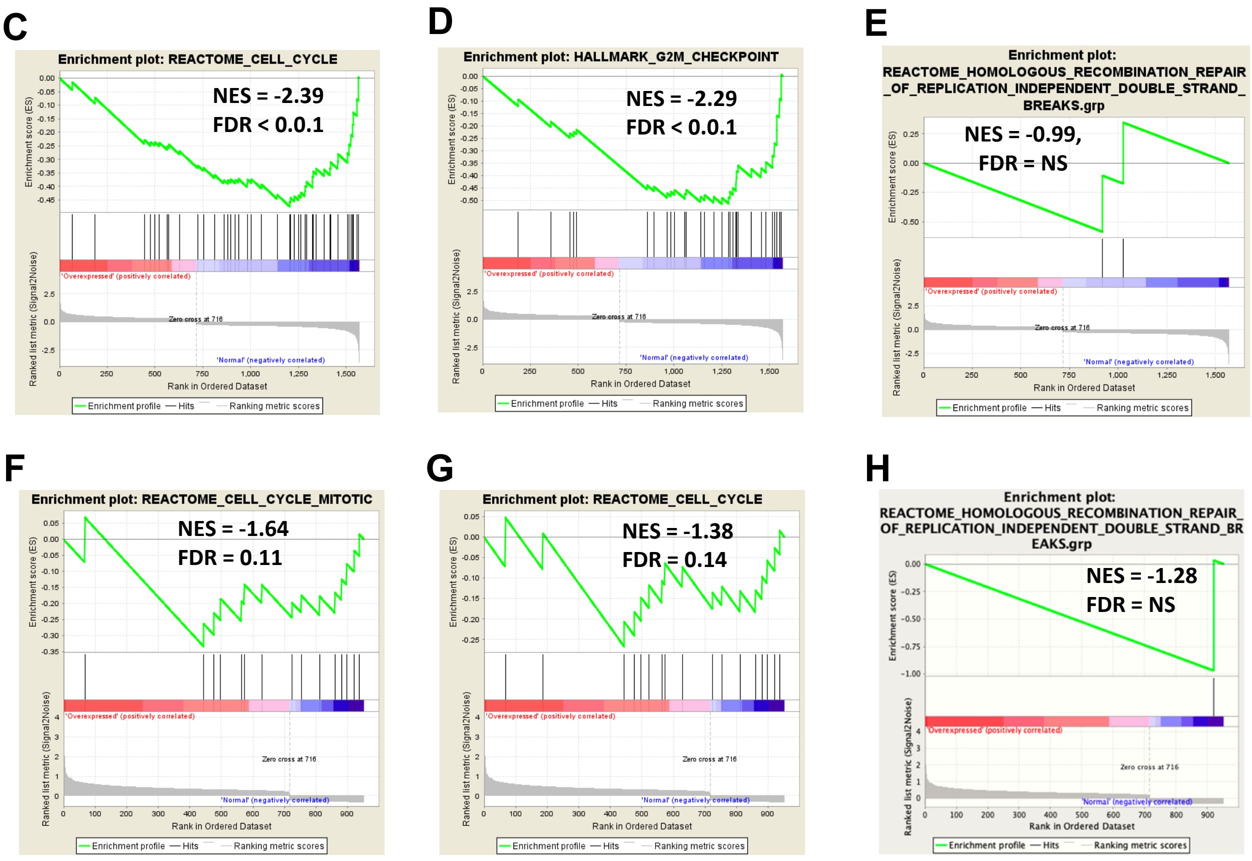


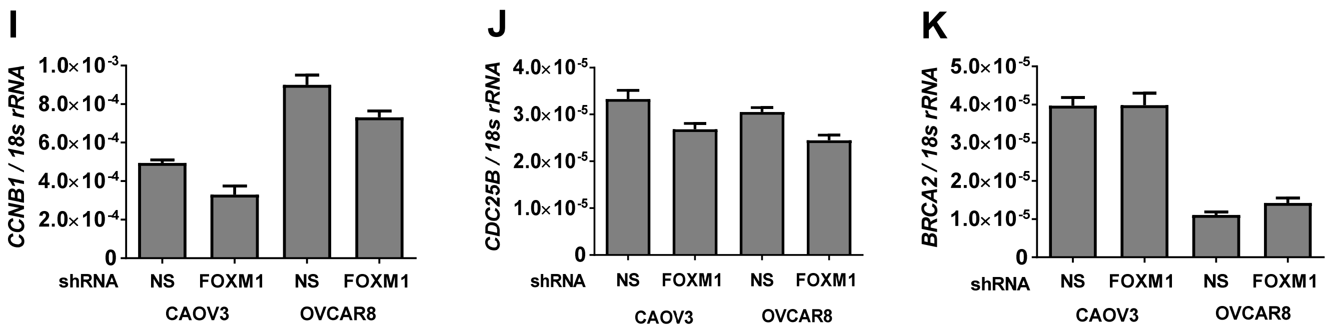

**Figure S14. Barger *et al*.**

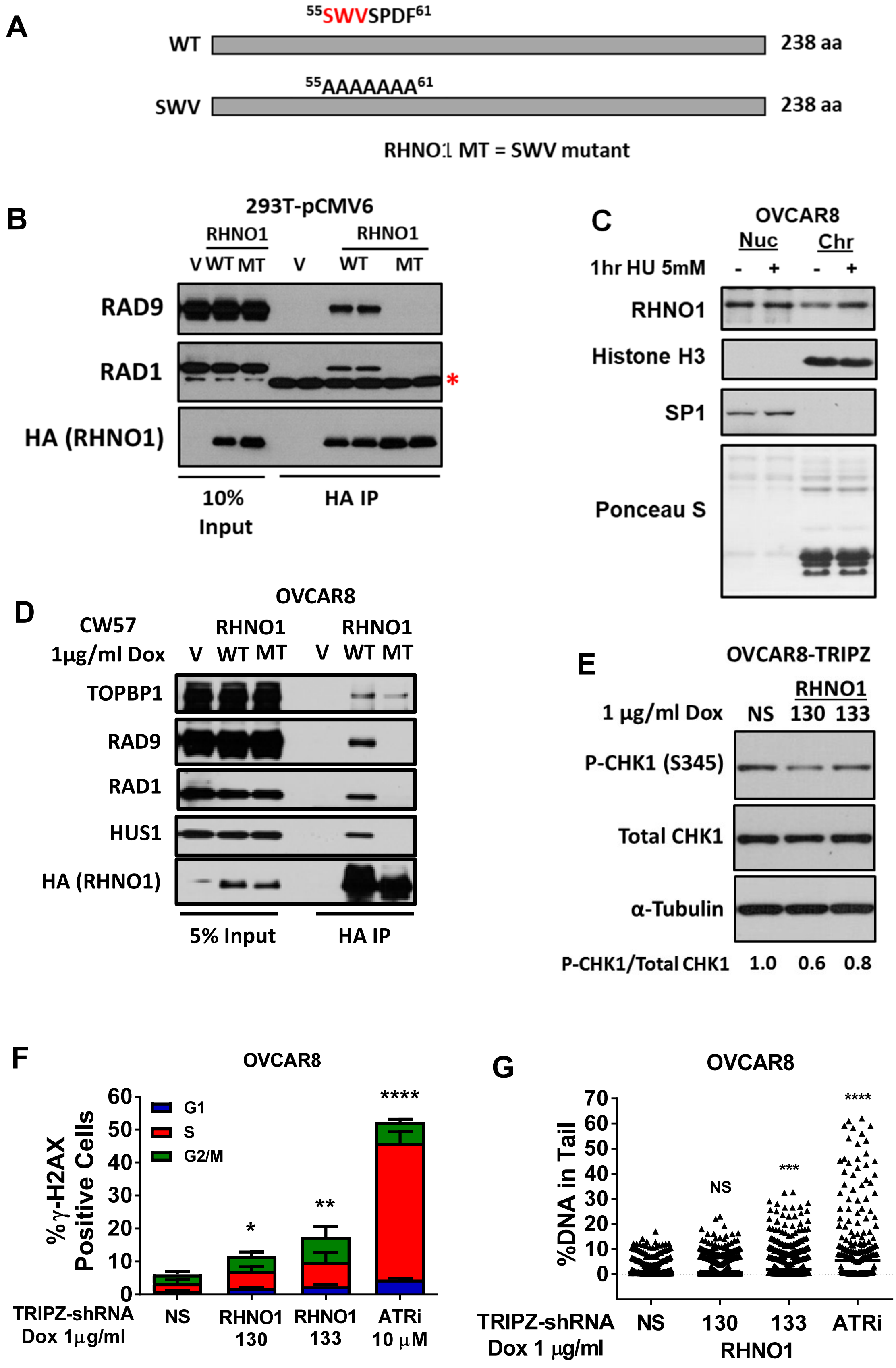

### Figure S15. Barger *et al*.

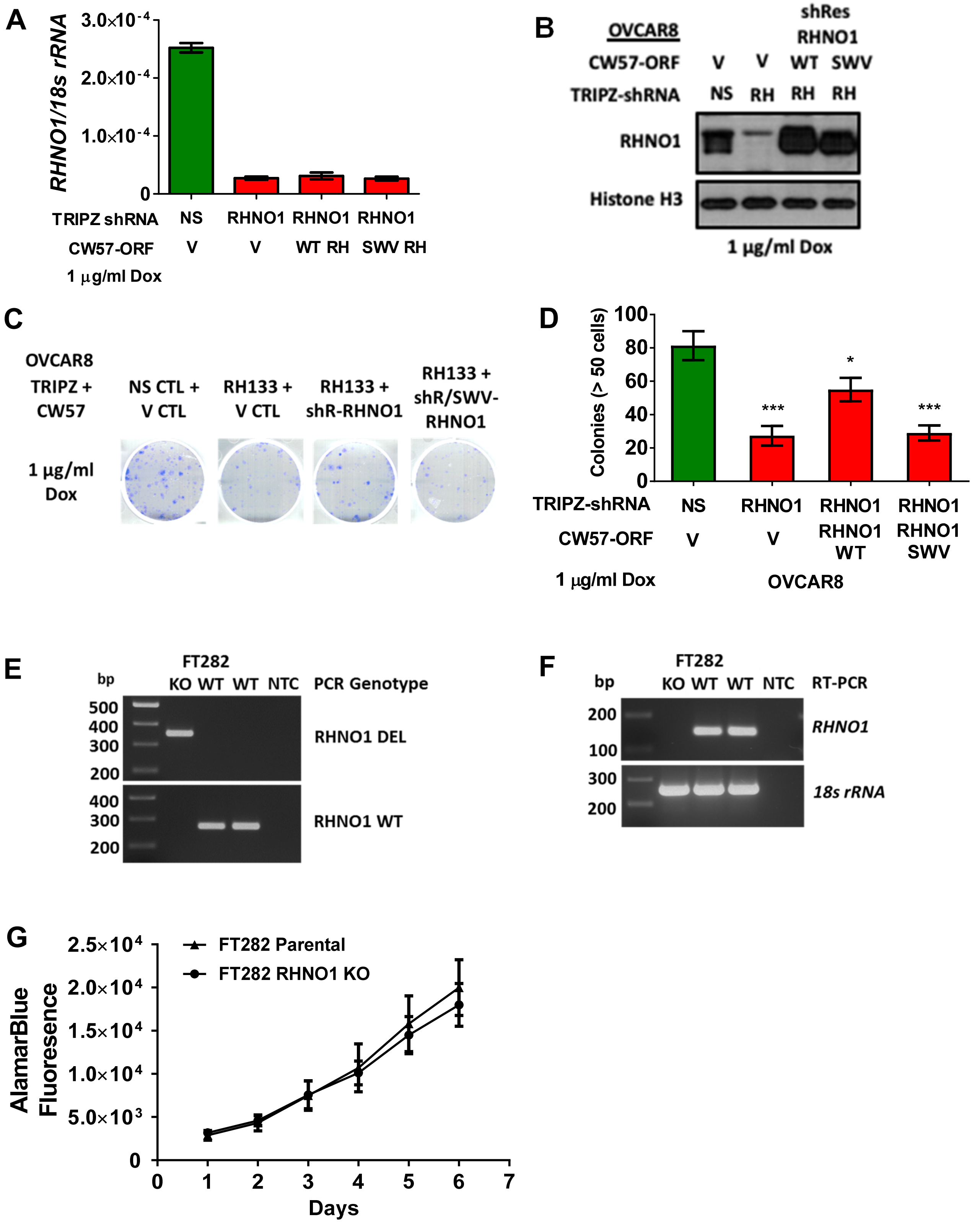
